# The characterisation of phytoene synthase-1 and 2, and 1-D-deoxy-xylulose 5-phosphate synthase genes from red chilli pepper (*Capsicum annuum*)

**DOI:** 10.1101/2023.01.25.525515

**Authors:** Harriet M. Berry, Nan Zhou, Daniel V. Rickett, Charles J. Baxter, Eugenia M. A. Enfissi, Paul D. Fraser

**Affiliations:** School of Biological Sciences, Royal Holloway, University of London, Egham, Surrey, TW20 OEX, UK; Syngenta, Jealott’s Hill International Research Centre, Bracknell, Berkshire, RG42 6EY, UK

**Keywords:** Isoprenoids, carotenoids, *Capsicum*, plastids, phytoene synthase (PSY), 1-*D*-deoxy-xylulose 5-phosphate synthase (DXS)

## Abstract

The red colouration of ripe chilli pepper fruit is due to the presence of the carotenoid capsanthin. 1-*D*-deoxy-xylulose 5-phosphate synthase (DXS) and phytoene synthase(s) (PSY) are influential biosynthetic steps in the carotenoid pathway. A panel of chilli accessions for varying fruit colour intensity revealed the correlation of carotenoid content with *PSY* and *DXS* transcript levels. The *PSY* and *DXS* genes were sequenced from high and low carotenoid genotypes to deduce potential allelic variation. *PSY*-1 and -2 showed tissue specific expression, with *PSY*-1and 2 expressed in chromoplast and chloroplast containing tissues, respectively. Protein modelling, phylogenetic analysis and transcription factor analysis was conducted to understand how the allelic variation identified could affect carotenoid biosynthesis and regulation, and subsequently colour intensity phenotype. A candidate mutation located in the transit peptide of the PSY-1 protein sequence was identified in the low colour intensity genotype. Within the promoter region of the *DXS* gene present in the high colour intensity genotype sequence variation was determined in the glyceraldehyde 3-phosphate binding motif (GAPF) of the promoter region. These data can be exploited to develop tools and resources for the breeding of high colour intensity chilli pepper fruit.

**Key policy highlights:** Using the molecular findings disclosed in this article the potential exists to develop molecular markers for the breeding varieties of chilli peppers with improved aesthetic consumer preference and nutritional attributes.

## Introduction

Chilli pepper is the world’s most widely grown and consumed spice, typically being used as a powder in cooking. The consumer associates the deep red colour of chilli pepper with flavour, maturity, and nutritional value. Therefore, the deep red colour of chilli powder is directly related to commodity price. The global production of fresh pepper reached 34.6 million tons and 3.5 million tons of dried pepper pods in 2014 (Qin et al., 2014).

Carotenoids are a distinctive class of natural pigments which cannot be synthesised in animals. In humans’ carotenoids are essential for optimal health and must be dietary acquired. For example, β-carotene (provitamin A) is essential for vision, vitamin A deficiency can lead to Xeropthalmia and potentially blindness (Krinsky and Johnson, 2005). The carotenoids lutein and zeaxanthin have also been linked to the reduction of macular degeneration (Congdon and West, 1999). Other health promoting properties of carotenoids are linked to their potent antioxidant properties, which can dissipate reactive oxygen species (ROS) resulting from cellular metabolism (Britton, 1995). In plants carotenoids act as ancillary pigments during photosynthesis and phytohormone precursors (Cazzonelli, 2011).

Biosynthetically, carotenoids are isoprenoids which are composed of eight isopentenyl diphosphate (IPP) isoprene units. Plastids synthesise IPP using the Methylerythritol 4-phosphate (MEP) pathway that utilises the glycolytic intermediates pyruvate and glyceraldehyde 3-phosphate (G3P). The first step in the MEP pathway, involves the formation of 1-*D*-deoxy-xylulose 5-phosphate (DXP) from pyruvate and G3P and is catalysed by the DXP synthase (DXS) (Figure. 1). In sweet pepper this enzyme has been referred to as *Capsicum* transketolase 2 or CapTKT2; however, in this study it will be referred to as DXS (Bouvier et al., 1998). DXS is considered to be the “rate limiting” step for isoprenoid biosynthesis in *Arabidopsis* and tomato (Lois et al., 2000). The outputs from the MEP pathway are IPP and Dimethylallyl pyrophosphate, these represent the universal C5 precursors from which all isoprenoids are formed. Geranylgeranyl pyrophosphate (GGPP) is a C20 prenyl lipid and represents the immediate isoprenoid derived precursor for carotenoids. GGPP is formed from three molecules of IPP and one molecule of DMAPP by the sequential action of geranyl pyrophosphate synthase (GPPS) and GGPP synthase (GGPPS) (Rodríguez-Concepción, 2004) (Figure 1). Phytoene synthase (PSY) is the first committed enzyme in the carotenoid biosynthesis pathway and encodes an enzyme which is responsible for the head to tail condensation of GGPP. The *Capsicum* carotenoid pathway has two branches, the α-branch, which is predominantly active in green tissue, and the β-branch, which is predominantly active during fruit ripening. The β-branch leads to the accumulation of capsanthin and its esters that are responsible for the fruits characteristic red colour (Figure 1).

**Figure 1.**
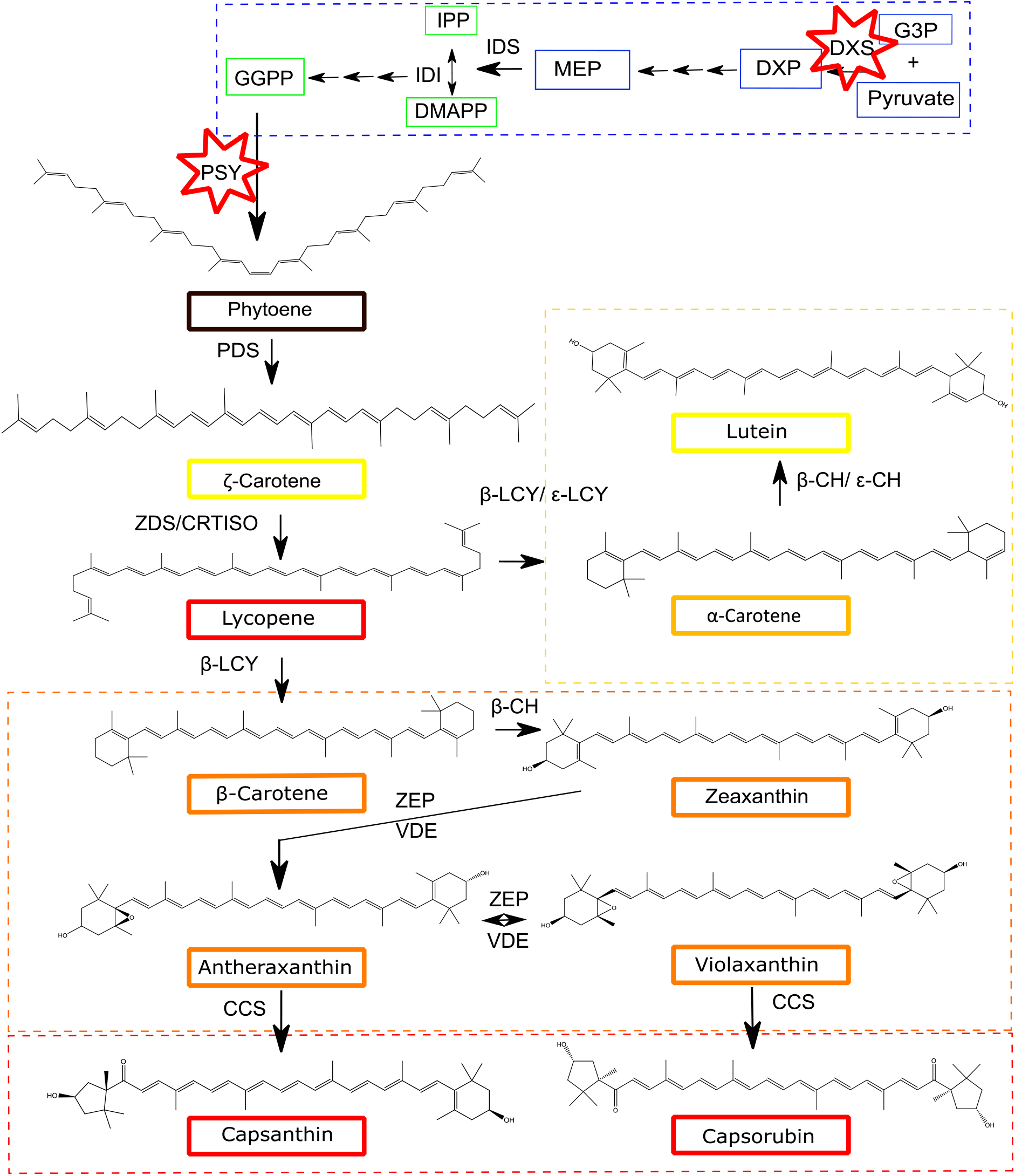
The carotenoid biosynthetic pathway in capsicum, using isopentenyl pyrophosphate (IPP) from the Methyl Erythritol Pathway (MEP). The positions of DXS and PSY in *Capsicum* are indicated with red stars. Pyruvate and G3P are used as precursors to the MEP pathway creating DXP. GGPP is the precursor for phytoene, the first committed step in carotenoid biosynthesis pathway. Key: yellow box, α-branch; orange box, β-branch; red box, xanthophylls in *Capsicum*. Abbreviations: MEP, 2-C-methyl-D-erythritol 4-phosphate; G3P, D-glyceraldehyde 3-phosphate; DXP, 1-deoxy-D-xylulose 5-phosphate; DXS, DXP synthase; MEcPP, 2-C-methyl-D-erythritol-2,4-cylcopyrophosphate; IPP, isopentyl disphosphate; DMAPP, dimethylallyl diphosphate; IDS, IPP/DMAPP synthase; IDI, IPP/DMAPP isomerase; GGPP, geranylgeranylpyrophosphate; PSY, phytoene synthase; PDS, phytoene desaturase; ZDS, ζ-carotene desaturase; CRTISO, carotenoid isomerase; LCY-β, lycopene β-cyclase; LCY-ε, lycopene ε-cyclase; β-CH, β-carotene hydroxylase; ε-CHY, ε-carotene hydroxylase; ZE, zeaxanthin epoxidase; VDE, violaxanthin de-epoxidase; CCS, capsanthin capsorubin synthase.

The present study builds on previous work characterising a fruit colour intensity panel in chilli pepper (Berry et al., 2019). These data have highlighted PSY and DXS as the key influential biosynthetic steps in the formation of carotenoids in chilli pepper fruit. The aim of this study was to further this hypothesis and characterise the *PSY* and *DXS* multigene families at the genomic level for putative SNPs responsible for altered fruit carotenoid content. Collectively, these data provided (i) fundamental insights into the regulation of carotenoid biosynthesis and potential tools and resources for explanation in breeding of high colour intensity chilli pepper fruit.

## Materials and Methods

### Germplasm and cultivation

The chilli pepper lines from this study were part of a colour intensity panel supplied by Syngenta (Berry et al., 2019). The plants were grown from seed in a temperature and light controlled glasshouse (25 °C day/15 °C night; 110 μmol m^−2^ s^−1^; 16 h/8 h light/dark cycle). The plants were tagged at anthesis as described in (Berry et al., 2019).

### Gene expression analysis by real time qPCR

RNA was extracted from three biological replicates; each biological replicate was an independent plant from the same genotype comprising three fruits. Frozen fresh pericarp tissue was ground to a fine powder using a tissue lyser (Qiagen) and extracted (50 mg) using RNeasy plant mini kit (Qiagen). Analysis of gene expression profiles was carried out using the SYBR green method described in (Enfissi et al., 2010). cDNA was generated using Illustra RT-PCR ready-to-go beads (GE Healthcare) in a two-step reaction. The reference gene, ATP synthase subunit alpha F1 complex, used in these studies was selected and tested using a geNorm kit (Primer design). Primer sequences for *PSY*-1 and -2 can be found in (Berry et al., 2019).

### Sequencing and modeling

The genes in this study were sequenced using primers designed to create overlapping fragments (Table S1-3). These primers were based on sequences available from the National Centre for Biotechnology (NCBI; http://www.ncbi.nlm.nih.gov) for sweet pepper (*PSY* and *DXS*) and tomato (*PSY-*2). Primers were designed using NCBI/Primer-BLAST and Primer3Plus (http://www.bioinformatics.nl.cgi-bin/primer3plus/primer3plus.cgi). Gene amplification was performed at 95°C (2min), then 30 cycles: 95°C (20 sec), 55-60°C (10 sec), and 72°C (20 sec). PCR reactions were carried out using Illustra puReTaq Ready-to-go Beads (GE Healthcare). Fragments were cloned using pCR 2.1 TOPO TA cloning kit and sequenced. The genes were cloned from genomic DNA to allow comparison of promoter and coding region. Construction of contigs was carried out using SeqMan Pro and Seqbuilder (Lasergene, DNAstar, UK). In silico 3D protein models was performed using SWISS-MODEL (http://swissmodel.Expasy.org) (Biasini et al., 2014).The 5′UTR from *PYS-*2 was annotated based on the tomato *PSY-*2 promoter (Giorio et al., 2008).Transit peptide prediction was carried out by Predotar (Small et al., 2004).

### Phylogenetic analysis

The sequences were mined from NCBI and Sol Genomics Network (https://solgenomics.net/). An uprooted neighbour-joining tree with 1000 bootstrap repetitions was constructed using the software MEGA-X (Kumar et al., 2018). The accession numbers and the phytoene synthase amino acid sequences used in construction of the phylogenetic tree can be found in Table S4.

### Transcription factor analysis

The *Nsite* programme was used to perform transcription factor analysis to allow identification of regulatory elements within the promoter region of the *PSY*-1, *PSY*-2 and *DXS* genes (http://www.softberry.com).

## Results and discussion

### The expression of *PSY* genes during chilli pepper fruit ripening

The chilli pepper genotypes designed R3 and R7 were confirmed as low and high carotenoid containing lines (Berry et al., 2019). Visual inspection indicated that the R3 line had an orange hue and contained approximately 3500 mg g^−1^DW, while the dark red R7 contained about 9000 mg g^−1^DW. This routinely correlated to the R7 line having a 2.5-fold significant increase in carotenoids compared to the R3 line.

PSY-1 has been shown to have a the most influence on the accumulation of carotenoids in tomato (Fraser et al., 2002), sweet pepper (Huh et al., 2001), and chilli pepper fruit (Berry et al., 2019). A second isoform of phytoene synthase (*PSY-*2) was identified and sequenced from Chilli pepper in this study in order to assess expression profiles and then gene characterisation.

The Chili pepper *PSY*-2 expression patterns were determined concurrently with *PSY*-1 (Figure. 2c and e). Although *PSY-*2 was expressed in ripe chilli fruit, the levels were very low when compared to *PSY-*1, for example a 2000-fold difference between *PSY-*1 and *PSY-*2 for the R7 line (Figure. 2e). This was double that of the R3 line when comparing *PSY-*2 to the *PSY-*1 expression peak (Figure. 2c). However, in both lines there was a similar level of expression of *PSY-*1 and *PSY-*2 in green tissues (e.g., leaf and green fruit) (Figure. 2b), with the onset of ripening, *PSY-*1 expression increased rapidly while *PSY-*2 decreased. Although low *PSY*-2 transcripts were still detectable in ripe fruit (Figure. 2d-f). These data infer that the product of *PSY*-2 is responsible for carotenoid formation in chloroplast containing tissues, while *PSY*-1 products initiate carotenoid biosynthesis in ripening of red chilli pepper fruit. The discovery that *PSY*-1 transcripts were over 2000-fold more abundant when compared to *PSY*-2 in ripe fruit from the high carotenoid Chilli line, which is represented a two-fold increase when compared to the low line, justified the detailed analysis of *PSY* at gene level. To date only one DXS associated with Chilli fruit has been annotated. DXS transcripts do increase during ripening in a similar pattern to *PSY*-1 but the peak considerably less pronounced (Berry et al., 2019).

**Figure. 2.**
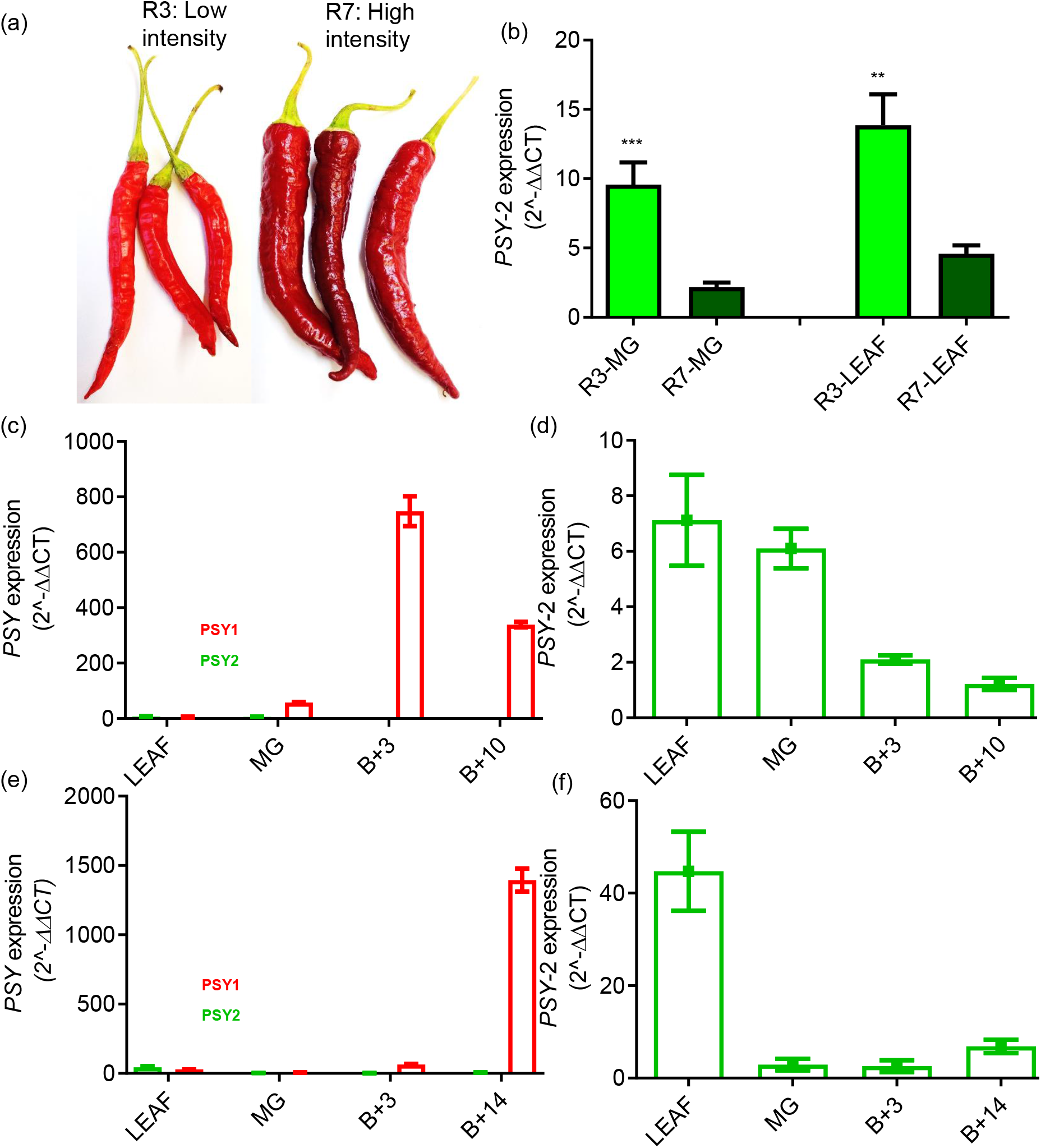
Comparison of *PSY*-1 and *PSY*-2 in chilli pepper lines differing in colour intensity phenotype. A low colour intensity line (R3) was compared to the high colour intensity line (R7). (a) Visual colour comparison for R3 and R7. (b) Comparison of the *PSY*-2 transcript levels in green tissue between lines. Comparison of *PSY*-1 and *PSY*-2 transcript levels in leaf and throughout ripening and development in fruit in R3 (c) and R7 (d). A refine profile for *PSY*-2 transcript levels in leaf and throughout fruit development and ripening in R3 (d) and R7 (f). Error bars represent ± SE (n=3). Student’s *t* test *** *P* ≤ 0.001.

The tissue specific expression of *PSY*-1 and 2 found in over Chilli pepper fruit development and ripening is similar to that found in tomato. In tomato detailed studies have determined PSY enzyme activity in a variety of mutants. These studies have confirmed the biosynthetic role of PSY-1 in the formation of carotenoids present in ripe fruit and PSY-2 in chloroplast containing tissues. In these studies it was concluded that PSY-2 does not contribute to the formation of carotenoids in ripe fruit (Fraser et al., 1999). Although recently it has been illustrated that under certain conditions PSY-2 has some contribution to carotenoid formation in ripe tomato fruit (Karniel et al., 2022).

### Phylogenetic analysis

An uprooted phylogenetic tree was constructed with 52 PSY sequences to investigate the phylogenetic relationship between the sequences obtained from R3 and R7 compared the other *Capsicum* varieties and many other higher plants (Figure. 3). The *Capsicum* PSY-1 accessions separated into two subgroups: A predominantly “orange” subgroup (subgroup 1 - containing one red variety) and a “red” subgroup (subgroup 2). The low intensity line was found in the “orange” subgroup and the high intensity line was found in the “red” subgroup. The “orange” and “red” descriptor presumably referring indirectly to carotenoid content Other subgroups were: *Solanaceae* PSY-1 (aubergine, tomato and potato), *Solanaceae* PSY-2 (*Capsicum*, aubergine, tomato and potato), eudicot PSY-1: subgroup 1 (loquat, cassava (PSY-1 and 2), *Arabidopsis*) and subgroup 2 (citrus, melon and watermelon), monocot PSY-1 subgroup 1 (saffron and banana) and subgroup 2 (maize, sorghum, rice, wheat), monocot PSY-2 (loquat, saffron, maize, rice, banana) and PSY-3 (*Capsicum*, saffron, maize, sorghum and rice, and citrus and melon PSY-2).

**Figure. 3.**
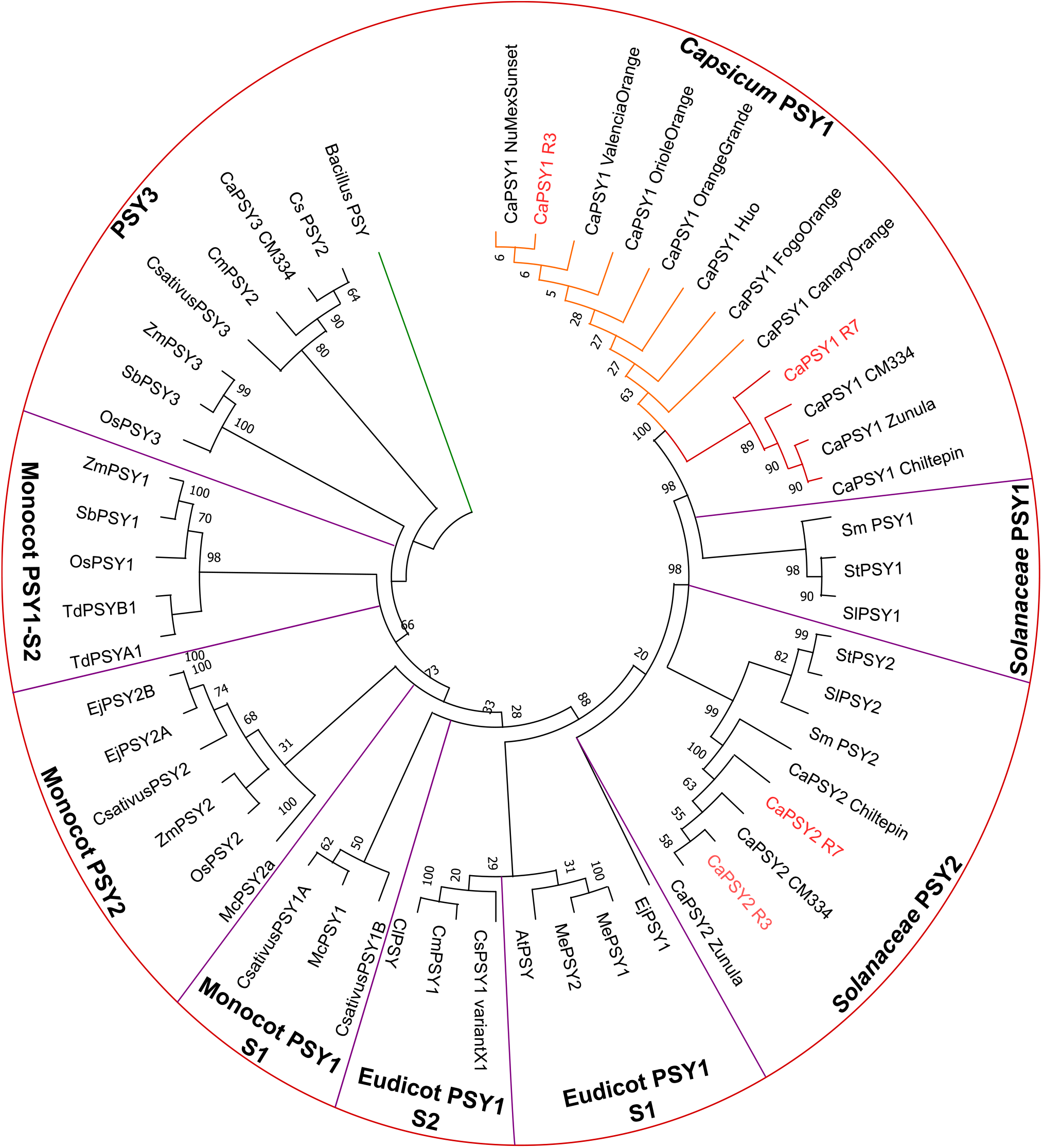
A phylogenetic tree of 52 PSY proteins from different plants. The phylogenetic tree of 52 PSY proteins was constructed using the software MEGA-X with neighbour-joining method to visualise the ancestry of R3 and R7 (red). A PSY from the *Bacillus* genus (WP_013058559.1) was used as an outgroup (green). *Capsicum* PSY1 contained two subgroups (orange subgroup, orange; red subgroup, red). Bootstrapping (1000 replicates) was used evaluate the degree of support for the grouping patterns displayed in phylogenetic tree. Accession numbers and amino acid sequences can be found in Supplementary material Table S4. Key: Ca, *Capsicum annuum*; At, *Arabidopsis thaliana*; Sm, *Solanum melongena*; Mc, *Musa cavendishii*; Me, *Manihot esculenta*; Cs, *Citrus sinensis*; Td, *Triticum durum*; Ej, *Eriobotrya japonica*; Cm, *Cucumis melo*; St, *Solanum tuberosum*; Os, *Oryza sativa*; Csativus, *Crocus sativus*; Sb, *Sorghum bicolor*; Sl, *Solanum lycopersicum*; Cl, *Citrullus lanatus*; Zm, *Zea mays*.

### Characterisation of the *PSY* and *DXS* genomic sequences and allelic variation

#### In silico phytoene synthase structure

The phytoene synthase enzyme belongs to a family of isoprenyl diphosphate synthases along with farnesylpyrophosphate synthase (FPS), geranylgeranyl pyrophospahte synthase (GGPS) and squalene synthase (SQS). Modelling of the protein structure of PSY was carried out based on previous models of dehydrosqualene synthase (Pandit et al., 2000) (Figure. 4a-b). The structure consisted of a central cavity which was surrounded by helices. The first layer of helices contains two aspartate-rich regions (DXXXD) which have been previously identified as essential for catalytic activity (Gu et al., 1998; Shumskaya et al., 2012) (Figure. 4d; black box). The negatively charged aspartate residues face into the cavity and are thought to bind the cofactor (Mg^+2^) and help stabilise the diphosphate groups. These regions were highly conserved between *PSY-*1 and *PSY*-2, from the R3 and R7 lines, as well as across other species which were compared in this study.

**Figure. 4.**
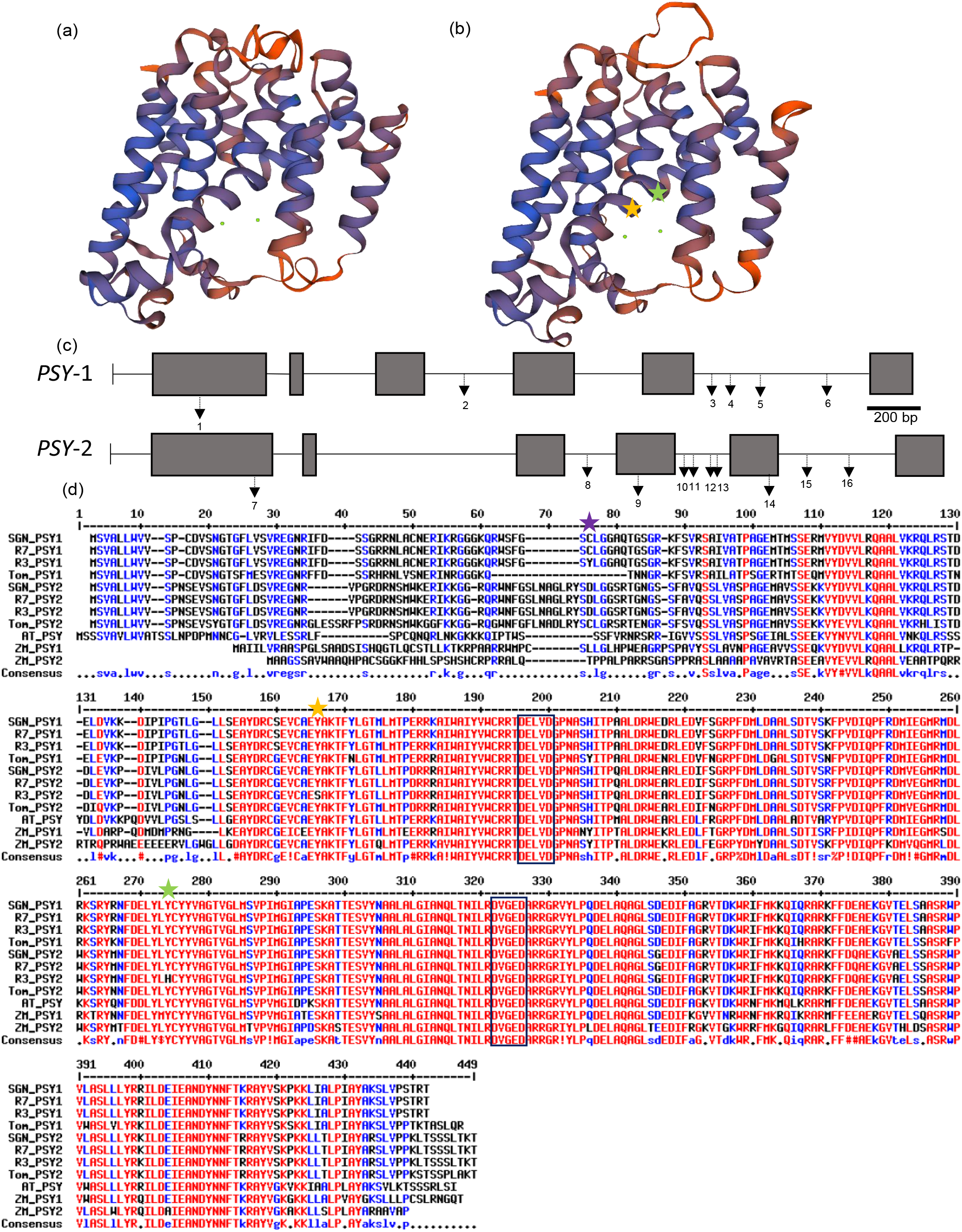
Sequencing of *PSY-*1 and *PSY*-2 genes. The *PSY-*1 and -2 gene sequences of chilli lines R3 and R7 were elucidated and the structure of the PSY1 protein was modelled using SWISSPROT. The phytoene synthase can bind two magnesium ion ligands (green dots in the central cavity) (a-b). (c) The gene structure and location of mutations present in *PSY*-1 and *PSY*-2: 1-R3 A^176^, 2-R3 G^1181^, 3-R3 G^2117^, 4-R3 C^2176^, 5-R3 A^2269^, 6-R3 A^2563^, 7-R3 C^446^, 8-R3 C^1632^, 9-R3 C^1816^, 10-R3 T^1988^, 11-R7 T^2051^ (insert), 12-R3 C^2126^, 13-R7 G^2160^ (insert), 14-R3 T^2362^, 15-R3 C^2505^ and 16-R3 T^2362^. Key: Black box, exon; black line, intron; arrows, mutations. (d) The coding regions of the *PSY*-1 and 2 genes in selected lines R3 and R7 were converted into an amino acid sequence and compared to other plant species available: SNG, Sol genomics network *Capsicum annuum* (PSY-1, CA04g04080; PSY-2, CA02g20350); Tom, *Solanum lycoperisum* (ABM45873); *Arabidopsis thaliana* (AT5G17230) and ZM, *Zea mays* (PSY-1, AAR08445; PSY-2, AAX13807). This revealed the presence of a base pair substitution in PSY-1 in R3 resulting in a cysteine replaced by a tyrosine (Tyr^59^) in the transit peptide (purple star) and two missense mutations present in PSY-2 in R3 (Ser^149^, orange star) and (His^257^, green star). The highly conserved regions present in the putative active site were conserved throughout (Black box) (Shumskaya et al., 2012). Alignment carried out using Multalin (Corbet, 1988).

#### Phytoene synthase-1 coding sequence

The coding sequences of *PSY-*1 from low (R3) and high (R7) colour intensity genotypes were determined. The *PSY-*1 gene contains 6 exons and 5 introns (Figure. 4c). The coding region is 1260bp, encoding a protein of 419 amino acids with a predicted molecular weight of 47.13 kDa. The gene is located on chromosome 4 of the pepper genome (Sol Genomics Network). The R3 amino acid sequence was identical to the GU085273 (Guzman et al., 2010) and X68017 accessions (Romer et al., 1993). R7 was similar, except for a tyrosine to cysteine change at residue 59 (Cys^59^) within the first exon of the coding region. This sequence was identical to CA04g04080 from the CM334 and the Zunla-1 genome (Sol genomics network). A chloroplast transit peptide was predicted with a probability score of 0.82. Although the tyrosine to cysteine substitution observed in R3 was located in the chloroplast transit peptide, and so unlikely to directly effect the activity of the enzyme. However it could potentially influence the translocation of the enzyme to the correct sub-plastid location, or other yet to be elucidated protein-protein interactions that confer optimal functionality. In addition, the amino substitutions were also identified in 6 orange varieties causing the R3 line to form a subgroup with these accessions when phylogenetic analysis was performed (Figure.3, *Capsicum* PSY-1 “orange”). There was one other “red” accession also found in this sub-group but the colour intensity phenotype relating to carotenoid content of this accession was unknown. These data imply that the low intensity phenotype line (R3) could perhaps be a dark orange phenotype, or a ‘false’ orange phenotype, whereby the orange colour is caused by a reduction in all carotenoids examined, including red coloured carotenoids. This profile is similar to accessions containing non-functional or impaired Capsorbin Capsanthin Synthase (CCS) enzymes. The orange colour phenotype occurs in these lines because of the accumulation of orange and yellow carotenoids. Therefore, the SNP present in the transit peptide could result in aberrant translocation of the PSY-1 protein within the plastid and thus consequently effecting activity. The identification of this SNP provides the potential opportunity to test and develop a molecular marker to screen (and potentially eliminate) low colour intensity phenotypes in pepper breeding.

#### Phytoene synthase-2 coding sequence

The sequence of chilli *PSY-*2 had not been published at the time the work was carried out. Therefore, based on homology with its tomato counterpart, the full sequence for the chilli *PSY-*2 was determined (Figure. 4). The Chilli *PSY*-2 contains 6 exons and 5 introns and is located on chromosome 2 (Sol Genomics Network) (Figure. 4c). The coding region is 1299 bp encoding a 438 amino acid protein with a predicted molecular weight of 48.33 kDa.

The amino acids sequences of the Chilli pepper PSY-2 showed greater similarity to the tomato PSY-2, than both Chilli and tomato PSY-1. Suggesting that the PSY-2 enzymes are more conserved between plant species that PSY-1. This is logical as PSY-1 is a paralog of PSY-2 derived from a gene duplication event (Giorio et al., 2008). The chilli PSY-1 was more similar to the tomato PSY-1, when compared to chilli PSY-2, likewise the chilli PSY-2 was more similar to the tomato PSY-2 than the chilli PSY-1 thus further supporting the identification of chilli *PSY-*2. *PSY-*1 and 2 have been sequenced in 94 *Capsicum* accessions a remarkably low incidence of mutations in the *PSY*-2 gene were found, further emphasising its essential role in plant development, when compared with the fruit tissue specificity of *PSY*-1 (Jeong et al., 2019). The gene alterations identified in this study were unique to the R3 and R7 lines and did not overlap with any of the mutations found by Jeong et al (2019).

A comparison of PSY-2 protein sequences between the R3 and R7 lines revealed two differences, which were unique to R3, even when comparison to other higher plants were made (Figure. 4d). These changes were a serine substitution for a tyrosine at residue 149 and histidine instead of tyrosine at residue 257 (Figure. 4b-c).

The amino acid substitutions identified in PSY-2 could potentially effect the activity of the enzyme. Both are located near to the central catalytic cavity. It was also found that the C-terminus of the PSY-2 protein is essential for the binding of two magnesium ion ligands. The absence of the last 60 amino acids results in the model only being able to bind one magnesium ion (Cao et al., 2019). When comparing the coding sequences of *PSY*-1 and *PSY*-2, an insert was located in the putative chloroplast transit peptide in PSY-2, which is also seen in tomato. This insert could potentially result in the translocation of these proteins to different sub-plastid locations (Gallagher, 2004). Previous studies determining the Capsicum PSY activity revealed that the sub-plastid location is fundamental to its optimal active state. This optimal state is found in the carotenoid biosynthesis membranes and stroma fractions derived from isolated chromoplasts (Berry et al., 2019).

Interestingly, the PSY1 maize mutant containing the Asn^168^ which gave rise to increased carotenoid production and fibrillar plastoglobuli phenotype was not present in the amino acid sequence of PSY1 in the chilli pepper lines selected. Even though *Capsicum* is known to accumulate fibrillar plastoglobuli (Deruère et al., 1994; Shumskaya et al., 2012). This suggests that the Asn^168^ amino acid is not responsible for the initiation of fibril formation in *Capsicum*. However, when the activity of the enzyme with the Asn^168^ mutant was removed, and so was no longer a highly expressed *PSY1*, fibrillar formation ceased to occur. Thus, suggesting transcriptional control leading to reduced activity, which indirectly affects coordinated fibrillar production.

#### DXS coding sequence

The *DXS* gene was amplified from both a low (R3) and high (R7) intensity genotype (Figure. 5). The coding regions are 4135 bp, containing 10 exons and 9 introns and the translated protein comprises 719 amino acids with a molecular weight of 77.60 kDa (Figure. 5c). A number of unique substitutions were identified on the R3 line (Val^77^, Ala^121^, and Cys^610^) compared to R7 and with the sweet pepper, tomato, *Arabidopsis* and maize sequences (Figure. 5d). The R7 contained two amino acid substitutions (Phe^251^ and Val^232^). The Val^232^ substitution was located within the conserved thiamine binding domain in the enzyme, as reported (Bouvier et al., 1998). This gave the DXS protein sequences of R3 and R7 99.3% similarity.

**Figure. 5.**
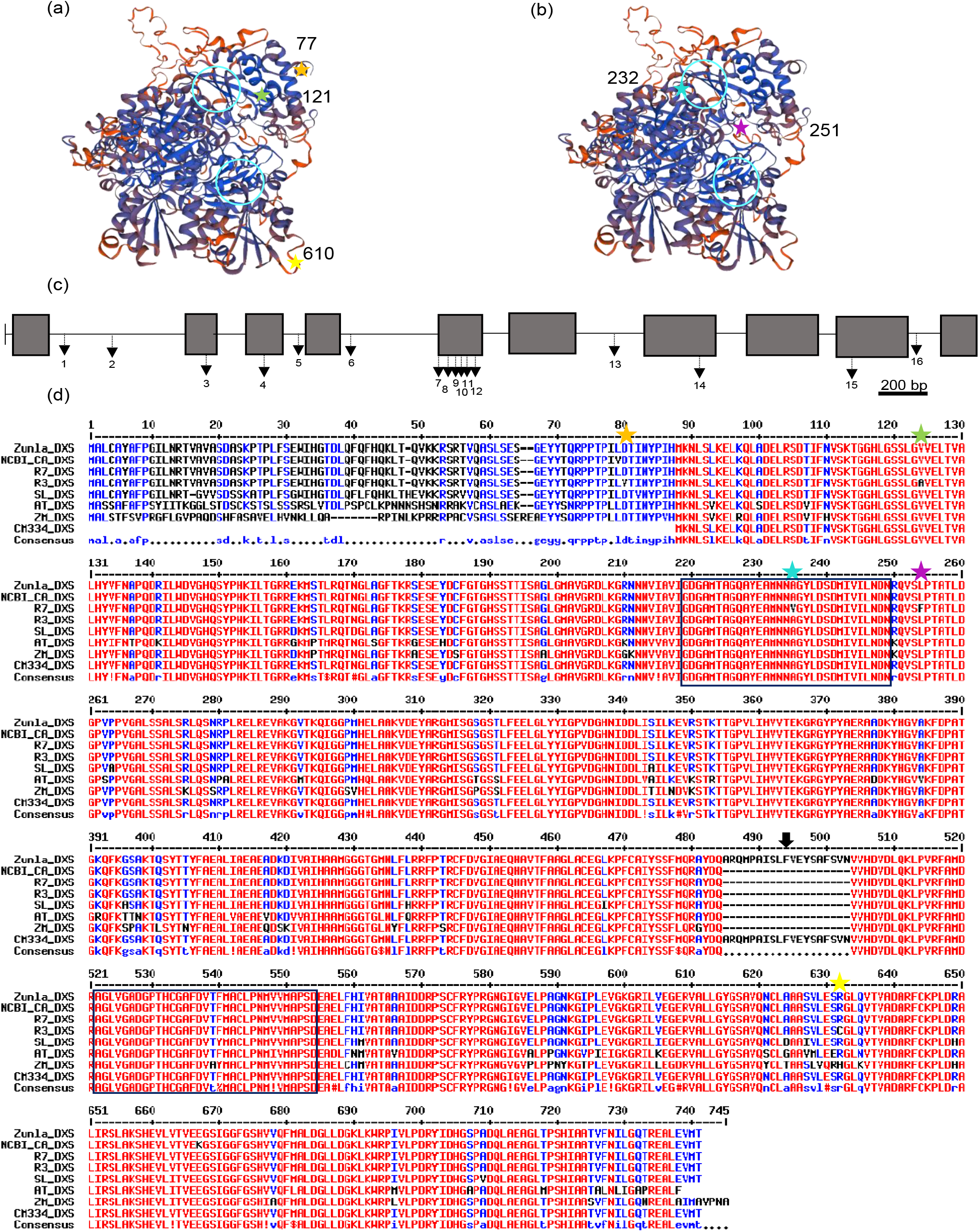
Sequencing of the 1-deoxy-D-xyulose 5-phospahte (*DXS*) gene. The *DXS* gene and promoter region was sequenced in a high (R7) and low intensity line (R3). 3D structures were constructed based on their amino acid sequence (a) R3 and (b) R7. The gene structure and location of mutations present in *DXS*: 1-R3 T^290^, 2-R3 AATTTGAA^380-388^ (insert), 3-R3 T^796^, 4-R3 C^1040^, 5-R7 G^1171^, 6-R7 ^AT1366-67^, 7-R7 T^1826^, 8-R7 C^1856^, 9-R7 T^1882^, 10-R7 T^1901^, 11-R7 G^1916^, 12-R7 T1940, 13-R7 G^2553^, 14-R7 G^2973^, 15-R3 T^3609^, 16-R3 T^3807^. Key: Black box, exon; black line, intron; arrows, mutations. The sequences were aligned with other published Capsicum sequences, as well as other higher plants to investigate conserved regions (d). These sequences were obtained from SGN: Zunla_DXS, Zunla-1 genome (Capana00g003194); CM334_DXS, CM334 genome (CA01g25140); SL_DXS, tomato genome ITAG4.0 (Solyc01g067890) and NCBI: CA, Capsicum annuum (CAA75778); AT, Arabidopsis thaliana (CP002688); ZM, Zea mays (EF507248). Key: Coloured stars and residue number represent amino acid substitutions unique to R3 (a) and R7 (b) and the light blue circles (3D structure) and black boxes (sequence) represent thiamine diphosphate binding site (first) and a conserved domain in thiamine-dependent enzymes (second) (Bouvier et al., 1998b). Black arrow corresponds to a potentially incorrectly annotated genome. Alignment carried out using Multalin.

Although it is unclear on the effect this has on the activity of the protein, it is unlikely to affect the binding drastically as both amino acids present (alanine and valine) have hydrophobic side chains and are similar in structure. It is also known that the Glu^449^ residue plays a crucial role in the interaction of the thiamine diphosphate molecule and the deprotonation of the thiozolium ring, and this amino acid is present in both lines investigated (Bouvier et al., 1998).

Alignment of the R3 and R7 DXS protein sequences with other *Capsicum* DXS sequences available from SGN and NCBI exposed some discrepancy between the annotations of intron-exon boundaries (Figure. 5d, black arrow). This was revealed by the presence of a 19bp insert in the amino acid sequence but no changes in the genomic nucleotide sequence, implying that perhaps the gene was annotated incorrectly. The incorrect annotation was present in the first sequenced pepper genome published (CM334), this annotation could have been extrapolated to other genome sequences after (Zunla-1 and Chiltepin). To further this the DXS sequence from the CM334 genome also had the N-terminus of the enzyme missing completely. Again, SNPs in the genomic nucleotide sequence which may have led to more incorrect annotation. This suggests that the quality of the sequencing of the *DXS* gene during the first hot pepper genome could be responsible for the incorrect annotation. Support for this finding is evident from the tomato gene, which has been more extensively manually annotated, and did not show this insert. These events highlight the value and integrity behind manual gene annotation and functional characterisation to complement the important information revealed from large genome sequencing projects.

### (i) Sequencing of the promoter regions revealed putative regulatory element motifs

#### *PSY-*1 *and* 2

The promoter regions of the *PSY-*1 gene were identical in the R3 and R7 lines and in other *Capsicum PSY-*1 promoter sequences (CA04g04080). *In Silico* transcription factor (TF) analysis scanned the promoter regions for regulatory elements bound by known transcription factors providing insight into the types of stimulus that result in their regulation (Shahmuradov and Solovyev, 2015).

This revealed a predominance of regulatory element (RE) motifs associated with light regulation and abiotic stresses, that are present in many phenylpropanoid/flavonoid and chlorophyll associated genes (Figure. S1-2, Table S5-6). In addition, possible master regulator CarG box motifs, which typically bind MADs box TFs, were found. The CarG box motif has been found to be involved in embryonic development, flowering, and cell wall modification (Hepworth et al., 2002). Analysis of the *PSY-*2 promoter revealed motifs associated with light regulation, MADs box regulation, phytohormone modulation (auxin and ABA) as well as chloroplast development. Despite the presence of similar motifs, alignments between *PSY-*1 and 2 revealed only 44.2% similarity between the two. *PSY-*2 promoter regions of R3 and R7 had a 99% similarity, but substitutions and a deletion were identified in R3. These sequence changes were present in regions associated with light and phytohormone effects. The differences in the REs found in *PSY-*1 and *PSY-*2 illustrate the more fundamental role played by *PSY-*2. Not only was this gene light and stress regulated, but phytohormone involvement postulated, all having roles in plant and chloroplast development. Interestingly, although *PSY*-1 and *PSY*-2 promoter regions shared little sequence similarity, similar REs suggest overlap in the roles of both these genes. For example, the light regulated *CAB* motif and the phytochrome a motif found in *PSY*-1 are involved in photosynthetic roles, and the *RIN* transcription factor binding motif found in *PSY*-2 is related to fruit ripening genes. This is also seen in tomato when the fruit specific *PSY*-1 gene is lost in the *r, r* mutant resulting in a reduction in carotenoid content in fruit and leaf (Fraser et al., 1999).

#### DXS

The *DXS* promoter regions contained putative regulatory motifs for light, stress, phytohormones (e.g. ABA and auxin) metabolism and a long GAAA repeat, typically associated with genes involved in primary and intermediary metabolism (Kim and Guiltinan, 1999; Yanagisawa, 2000) (Figure. 5a-e, Table S7). The R3 and R7 *DXS* promoter regions had a 99% similarity, but a key difference was in differences in putative REs, which could be responsible for altering the regulation of *DXS*. A G-box motif found to bind glyceraldehyde 3-phosphate (GAP) binding factor (GAPF), which is also found in the promoter region of the glyceraldehyde 3-phosphate dehydrogenase (*GapA*), was identified in R3, but not in R7, due to a thymine to adenine substitution (Jeong and Shih, 2003) (Figure. 6b). There were three other substitutions identified in the promoter region, but they were not located in putative REs.

**Figure. 6.**
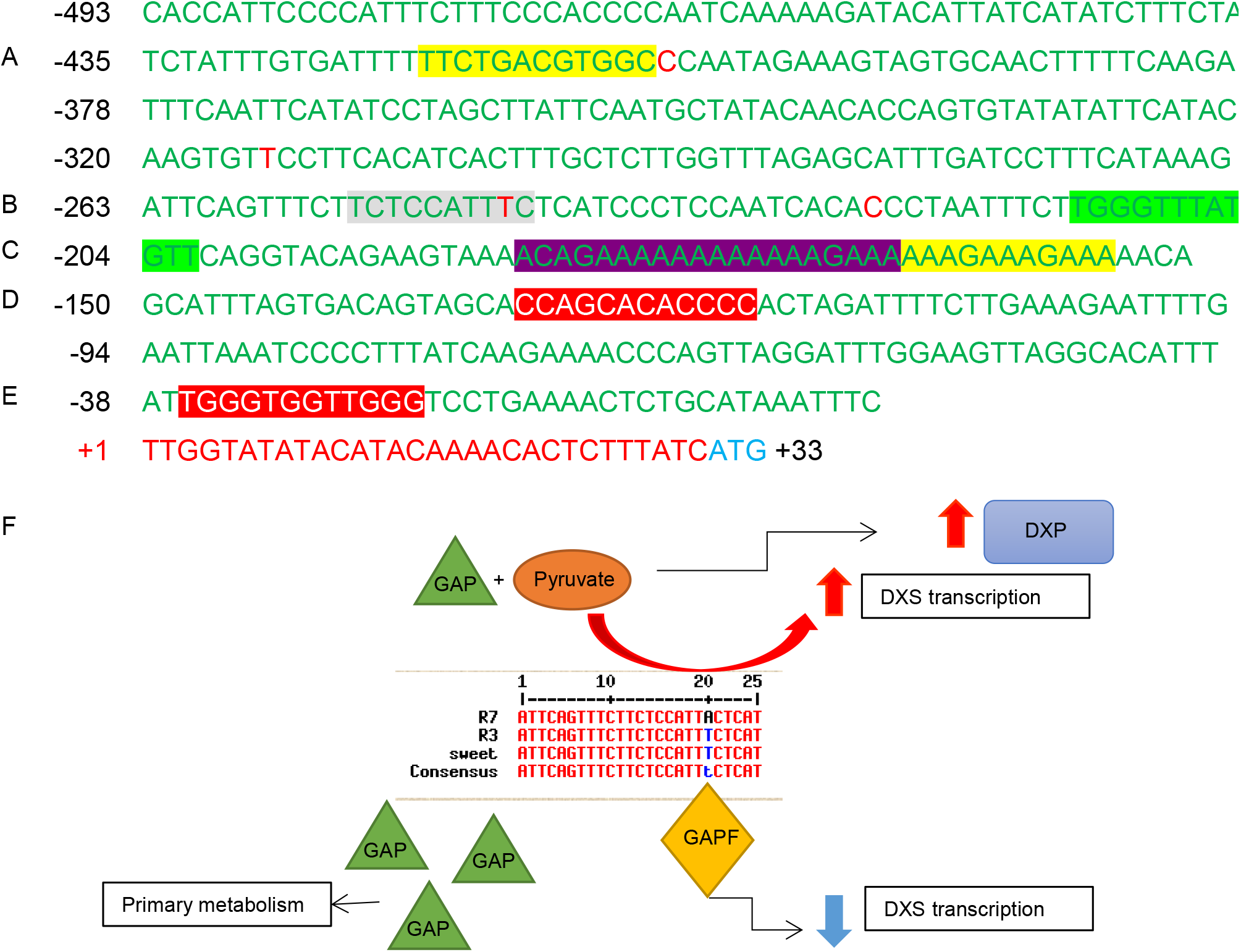
Putative regulatory elements present in the *DXS* promoter. Transcription factor analysis was carried out on the promoter region of DXS in R3 and R7 and putative regulatory elements (REs) were elucidated. The highlighted sequences correspond to Table S7. Yellow represents light REs (**A** and **C** (II)); light green represents plastid specific RE (**B** (II)); purple represents REs found in the promoters of genes involved in primary metabolism (**C** (I)) and red represents REs found in phenylpropanoid biosynthesis genes (**D**-**E**). The red bps indicate the differences between the R3 and R7 line and grey represents a G-box RE that is only present in R3. This above sequence is taken from the R3 line. (**F**) Diagram of proposed DXS promoter regulation whereby an amino substitution within a G-box motif could cause the R7 line to be expressed differently to R3.

The *DXS* gene displayed REs related to light, stress, and hormones, similar to *PSY-*2, in addition to REs related to sugar metabolism; thus, highlighting the role of DXS in channelling the flow of carbon from sugar metabolism into the isoprenoid pathway during the chloroplast to chromoplast differentiation. The difference in putative REs illustrates the presence of complex signalling pathways which synchronise fruit development and ripening, with the accumulation of carotenoids at the onset of ripening, as well as in response to stress. The REs also demonstrated the different roles between these genes, whereby *DXS* and *PSY-*2 have more links to primary metabolism and plant development, when compared to *PSY-*1, which is a fruit-specific gene, expressed at the onset of ripening (Figure. 6).

Sequencing of the *PSY-*1 and *DXS* genes in high (R7) and low (R3) intensity lines provided further evidence for the possibility of the underlying role *DXS* plays in the high colour intensity phenotype. In the high and low lines, the *PSY-*1 gene and promoter region are highly conserved between both lines, as well as with others characterised from the *Capsicum* genus (Figure. S1-2 and Table S5-6). However, the *DXS* gene was found to have 5 amino acid substitutions, and additionally, the *DXS* gene showed a mutation in a putative GAPF binding motif in R7, when compared to R3 (Figure. 6b). This motif was also present in the promoter region of *GapA*. These two genes share the same substrate, G3P, suggesting that a TF which modulates substrate flow from primary into secondary metabolism could be present. The flow of carbon is either needed for the plant to produce energy through glycolysis (primary) or to divert the carbon into the carotenoid pathway (secondary). However, the R7 line, is missing this motif, which provides a testable hypothesis to assess the interactions between primary metabolism and carotenoid formation. Interestingly, the metabolite profiles these lines did show reveal changes in those intermediary metabolites associated with changes in carotenoid and volatile formation (Berry et al., 2019, 2021).

## Concluding remarks

The present study builds on Berry et al (2019) that identified quantitative differential expression of carotenoid biosynthesis associated with altered carotenoid/colour intensity. Allelic variation in both *PSY*-1 and *DXS* genes associated with these altered contents have been identified providing a new testable hypothesis for the development of gene specific markers underlying colour intensity in chilli pepper fruit and potentially for other Solanaceae. Further flux analysis could shed more light on the connections with intermediary metabolism and the transcriptional regulation of carotenoid biosynthesis.

## Accession Numbers

Sequence data from this article can be found in EMBL/GenBank data libraries under accession numbers: KX588713-PSY1-R3, KX588714-PSY1-R7, KX588715-PSY2-R3, KX588716-PSY2-R7, KX588717-DXS-R3 and KX588718-DXS-R7.

## Author contributions

PDF secured the funding for the project. PDF, HMB, DVR and CJB conceived and designed research. HMB carried out gene expression analysis, sequencing and modelling. NZ carried out transcription factor analysis. HB wrote the original manuscript draft. PDF and GE edited the manuscript. All authors read and approved the manuscript.

## Acknowledgements

Financial support for this research was provided by BBRSC Syngenta Ltd. iCASE award BB/1015590/1 to PDF and GE.

