## Supplementary figures for "The characterisation of phytoene synthase-1 and 2, and 1-D-deoxy-xylulose 5-phosphate synthase genes from red chilli pepper (*Capsicum annuum*)"

### Supplemental material – Figures

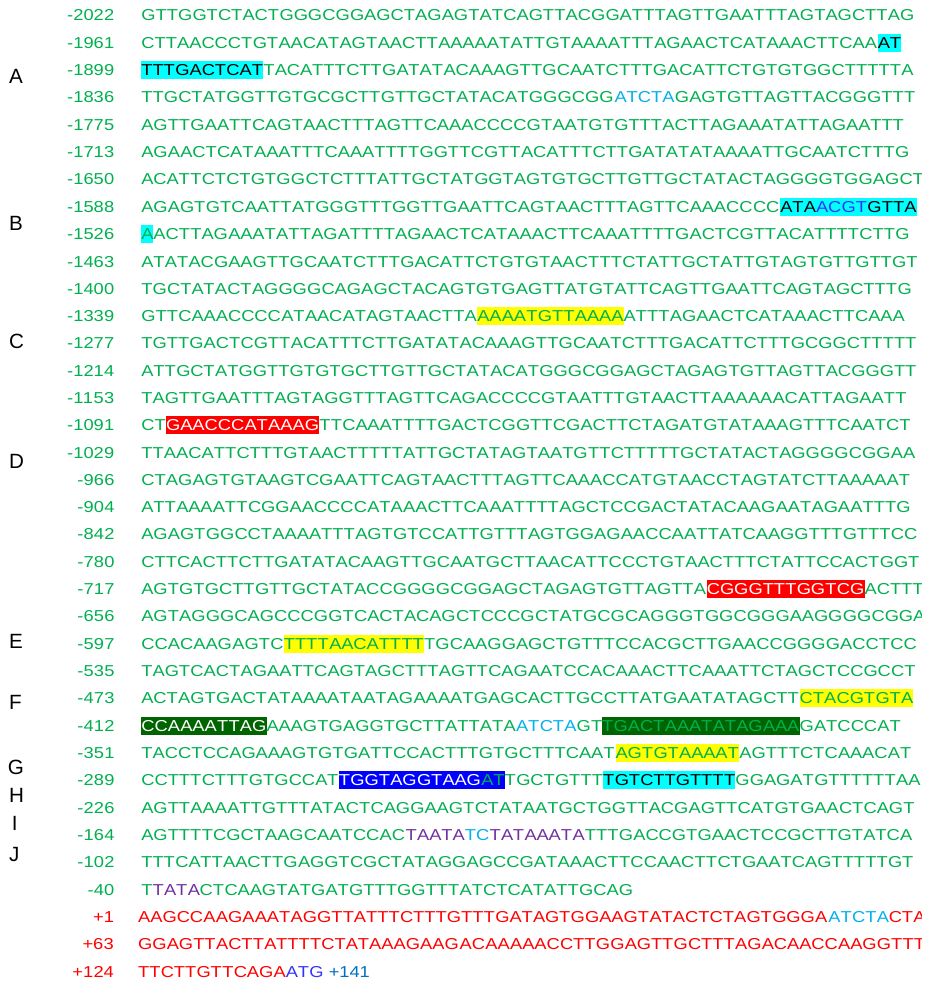

**Fig. S1. Putative regulatory elements present in the PSY-1 promoter.**

*In silico* transcription factor analysis was carried on the promoter of the *PSY-*1 gene and putative regulatory elements (REs) were elucidated. The blue ATG represents the start of the coding region, the red nucleotides represent the 5’UTR, and the green nucleotides represent the promoter region. The yellow highlighted sequences are light-regulated elements, the red highlighted sequences represent REs found in the phenylpropanoid biosynthesis pathway, light blue highlighted regions represent REs which are activated in response to stress, dark green represents CarG motifs, and dark blue represents a region which has the most homology with other promoters and possible key regulatory site. The light blue ATCTA motifs are also found in PSY of Arabidopsis (Welsch et al., 2003). Green indicates a possible TATA-rich motif as well as the purple TATA box. The letters **A**-**K** are described in Table S1.

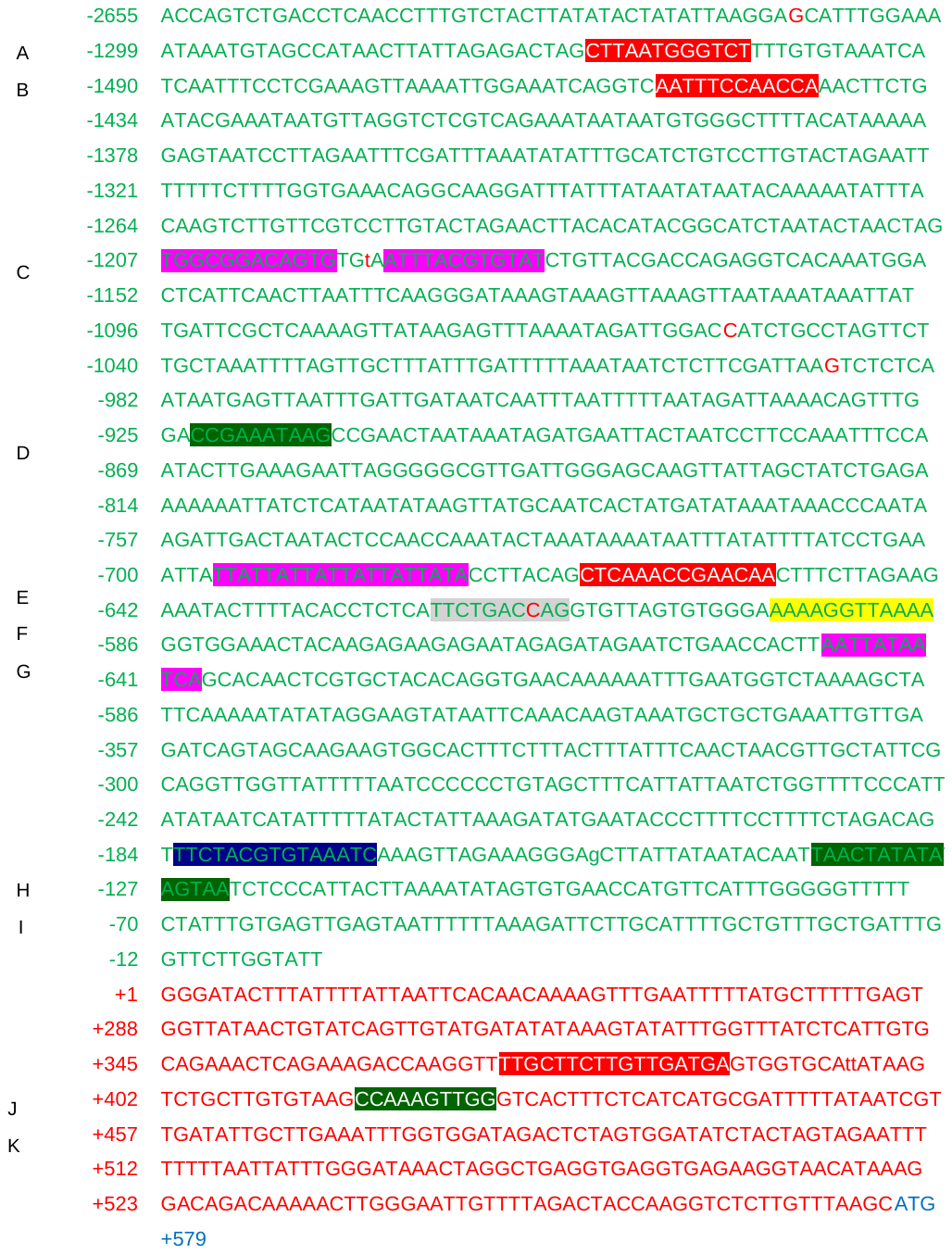

**Fig. S2. Putative regulatory elements present in the PSY-2 promoter.**

In silico transcription factor analysis was carried out on the PSY-2 promoter regions of R3 and R7 to elucidate putative regulatory elements (REs). The blue ATG represents the start of the coding region, the red nucleotides represent the 5’UTR and the green nucleotides represent the promoter region. The red highlighted regions represent REs found in phenylpropanoid biosynthesis related genes; the pink regions are REs found to be hormonally regulated; the light blue regions represent stress regulated REs; the dark green regions represent CarG box REs and the dark blue region is an element which has the most homology with other promoters and a possible key regulatory site The above sequence is the R7 promoter region and the differences between R3 and R7 are displayed as red characters and the grey region corresponds to a RE only present in R7. See Table S2 for description of REs.
