## Supplementary tables for "The characterisation of phytoene synthase-1 and 2, and 1-D-deoxy-xylulose 5-phosphate synthase genes from red chilli pepper (*Capsicum annuum*)"

### **Supplemental Material – Tables**

**Table S1. Primers designed for *PSY-1* sequencing.**

| **Name** | **Forward** | **Reverse** | **Annealing Tm** |
| --- | --- | --- | --- |
| P1 | TCTTTGACATTCTTTGCGGC | ACACTTTCTGGAGGTAATGGGA | 50 |
| P2 | GTGTGTGTTGGTCTACTGGG | AGGCCACTCTCAAATTCTATTCT | 53 |
| P3 | TCTTTGACATTCTTTGCGGC | AGGCCACTCTCAAATTCTATTCT | 53 |
| P4 | CACTACAGCTCCCGCTATGC | ACACTTTCTGGAGGTAATGGGA | 53 |
| P5 | TGCAGAGTACGCAAAGACGT | AGCTGATTTGCGATCCCCAA | 53 |
| P6 | CTCAGGAAGTCTATAATGCTGGT | CCCAAGCAAGAACCAAAACTCC | 53 |
| P7 | GATGCTGCTTTGTCCGACAC | AGCAAATGACGACACCCAGT | 53 |
| P8 | CTCAGGAAGTCTATAATGCTGGT | ATGTCAAATGGCCGTCCACT | 53 |
| P9 | GGCAGGTCTATCCGACGAAG | AGCAAATGACGACACCCAGT | 53 |
| P10 | GGCAGGTCTATCCGACGAAG | CCAGTTTTCGTGCCCATAGC | 53 |
| P11 | TACCGCAGGATACTGGACGA | CCAGTTTTCGTGCCCATAGC | 53 |

**Table S2. Primers designed for *PSY-2* sequencing.**

| **Name** | **Forward** | **Reverse** | **Annealing Tm** |
| --- | --- | --- | --- |
| F1 | ACCAGTCTGACCTCAACCTT | CCAACCTGCGAATAGCAACG | 56 |
| F2 | GGTGGAAACTACAAGAGAAGAGA | CCAAGAACCAAATCAGCAAACAG | 57 |
| F3 | ACCCTTTTCCTTTTCTAGACAGT | ACATCACCGTATATTGCCCAG | 56 |
| F4 | ACCCTGCTAATGACTCCAGAC | CCGCCCGCTGAAAATATCTT | 56 |
| F5 | TCCTGTTTCGTTTGGCAGTC | GCAAATATGTCTTCGCCGGA | 56 |
| F6 | CAGGGCTCTCCGGCGAAG | AACAGCAACGATGCCAACAC | 56 |

**Table S3. Primers designed for *DXS* sequencing.**

| **Name** | **Forward** | **Reverse** | **Annealing Tm** |
| --- | --- | --- | --- |
| D1 | ACACCACTTACCATTGAGGGG | CACTGCCACTGTCCTGTTCA | 55 |
| D2 | AGTGACAGTAGCACCAGCAC | AGAACTACATGTGGGAGAACTGT | 55 |
| D3 | TTGTAGAATGCTCATGTTGAACCA | TCCGACCGCTTAGTAAATCCA | 55 |
| D4 | TGTTGCAGTCATATCCTCACA | TGCGACTTCTCTTAGTTCTCTGA | 55 |
| D5 | CCAGTTCCTCCTGTTGGAGC | TTTAAGGCGGTCTAGTGGCG | 55 |
| D6 | AGGTCCTGTACTGATCCATGT | TGCACAGAAAGGTTTGAGGC | 55 |
| D7 | AGCAGATAAAGACATTGTTGCAA | ACTGCTGAGCCGTATCCCAA | 55 |
| D8 | ACTTTCATGGCATGTCTCCC | ATTCTTTTACAGTTCTTGCATCT | 55 |

**Table S4. Amino acid sequences and accessions used in phylogenetic analysis.** Corresponds to Fig. 3**.**

| **Plant family and accession** | **Amino acid sequence** |
| --- | --- |
| *Capsicum annuum*  Phytoene synthase-1  R3  (KX588713) | MSVALLWVVSPCDVSNGTGFLVSVREGNRIFDSSGRRNLACNERIKRGGGKQRWSFGSYLGGAQTGSGRKFSVRSAIVATPAGEMTMSSERMVYDVVLRQAALVKRQLRSTDELDVKKDIPIPGTLGLLSEAYDRCSEVCAEYAKTFYLGTMLMTPERRKAIWAIYVWCRRTDELVDGPNASHITPAALDRWEDRLEDVFSGRPFDMLDAALSDTVSKFPVDIQPFRDMIEGMRMDLRKSRYRNFDELYLYCYYVAGTVGLMSVPIMGIAPESKATTESVYNAALALGIANQLTNILRDVGEDARRGRVYLPQDELAQAGLSDEDIFAGRVTDKWRIFMKKQIQRARKFFDEAEKGVTELSAASRWPVLASLLLYRRILDEIEANDYNNFTKRAYVSKPKKLIALPIAYAKSLVPSTRT |
| *Capsicum annuum* Phytoene synthase-1  R7  (KX588714) | MSVALLWVVSPCDVSNGTGFLVSVREGNRIFDSSGRRNLACNERIKRGGGKQRWSFGSCLGGAQTGSGRKFSVRSAIVATPAGEMTMSSERMVYDVVLRQAALVKRQLRSTDELDVKKDIPIPGTLGLLSEAYDRCSEVCAEYAKTFYLGTMLMTPERRKAIWAIYVWCRRTDELVDGPNASHITPAALDRWEDRLEDVFSGRPFDMLDAALSDTVSKFPVDIQPFRDMIEGMRMDLRKSRYRNFDELYLYCYYVAGTVGLMSVPIMGIAPESKATTESVYNAALALGIANQLTNILRDVGEDARRGRVYLPQDELAQAGLSDEDIFAGRVTDKWRIFMKKQIQRARKFFDEAEKGVTELSAASRWPVLASLLLYRRILDEIEANDYNNFTKRAYVSKPKKLIALPIAYAKSLVPSTRT |
| *Capsicum annuum* Phytoene synthase-1  CanaryOrange  (GU085277) | MSVALLWVVSPCDVSNGTGFLVSVREGNRIFDSSGRRNLACNERIKRGGGKQRWSFGSYLGGAQTGSGRKFSVRSAIVATPAGEMTMSSERMVYDVVLRQAALVKRQLRSTDELDVKKDIPIPGTLGLLSEAYDRCSEVCAEYAKTFYLGTMLMTPERRKAIWAIYVWCRRTDELVDGPNASHITPAALDRWEDRLEDVFSGRPFDMLDAALSDTVSKFPVDIQPFRDMIEGMRMDLRKSRYRNFDELYLYCYYVAGTVGLMSVPIMGIAPESKATTESVYNAALALGIANQLTNILRDVGEDARRGRVYLPQDELAQAGLSDEDIFAGRVTDKWRIFMKKQIQRARKFFDEAEKGVTELSAASRWPVLASLLLYRRILDEIEANDYNNFTKRAYVSKPKKLIALPIAYAKSLVPSTRT |
| *Capsicum annuum* Phytoene synthase-1  FogoOrange  (GU085275) | MSVALLWVVSPCDVSNGTGFLVSVREGNRIFDSSGRRNLACNERIKRGGGKQRWSFGSYLGGAQTGSGRKFSVRSAIVATPAGEMTMSSERMVYDVVLRQAALVKRQLRSTDELDVKKDIPIPGTLGLLSEAYDRCSEVCAEYAKTFYLGTMLMTPERRKAIWAIYVWCRRTDELVDGPNASHITPAALDRWEDRLEDVFSGRPFDMLDAALSDTVSKFPVDIQPFRDMIEGMRMDLRKSRYRNFDELYLYCYYVAGTVGLMSVPIMGIAPESKATTESVYNAALALGIANQLTNILRDVGEDARRGRVYLPQDELAQAGLSDEDIFAGRVTDKWRIFMKKQIQRARKFFDEAEKGVTELSAASRWPVLASLLLYRRILDEIEANDYNNFTKRAYVSKPKKLIALPIAYAKSLVPSTRT |
| *Capsicum annuum* Phytoene synthase-1  Huo  (ACE78189) | MSVALLWVVSPCDVSNGTGFLVSVREGNRIFDSSGRRNLACNERIKRGGGKQRWSFGSYLGGAQTGSGRKFSVRSAIVATPAGEMTMSSERMVYDVVLRQAALVKRQLRSTDELDVKKDIPIPGTLGLLSEAYDRCSEVCAEYAKTFYLGTMLMTPERRKAIWAIYVWCRRTDELVDGPNASHITPAALDRWEDRLEDVFSGRPFDMLDAALSDTVSKFPVDIQPFRDMIEGMRMDLRKSRYRNFDELYLYCYYVAGTVGLMSVPIMGIAPESKATTESVYNAALALGIANQLTNILRDVGEDARRGRVYLPQDELAQAGLSDEDIFAGRVTDKWRIFMKKQIQRARKFFDEAEKGVTELSAASRWPVLASLLLYRRILDEIEANDYNNFTKRAYVSKPKKLIALPIAYAKSLVPSTRT |
| *Capsicum annuum* Phytoene synthase-1  NuMexSunsetOrange  (GU085274) | MSVALLWVVSPCDVSNGTGFLVSVREGNRIFDSSGRRNLACNERIKRGGGKQRWSFGSYLGGAQTGSGRKFSVRSAIVATPAGEMTMSSERMVYDVVLRQAALVKRQLRSTDELDVKKDIPIPGTLGLLSEAYDRCSEVCAEYAKTFYLGTMLMTPERRKAIWAIYVWCRRTDELVDGPNASHITPAALDRWEDRLEDVFSGRPFDMLDAALSDTVSKFPVDIQPFRDMIEGMRMDLRKSRYRNFDELYLYCYYVAGTVGLMSVPIMGIAPESKATTESVYNAALALGIANQLTNILRDVGEDARRGRVYLPQDELAQAGLSDEDIFAGRVTDKWRIFMKKQIQRARKFFDEAEKGVTELSAASRWPVLASLLLYRRILDEIEANDYNNFTKRAYVSKPKKLIALPIAYAKSLVPSTR |
| *Capsicum annuum* Phytoene synthase-1  OrangeGrande  (GU085276) | MSVALLWVVSPCDVSNGTGFLVSVREGNRIFDSSGRRNLACNERIKRGGGKQRWSFGSYLGGAQTGSGRKFSVRSAIVATPAGEMTMSSERMVYDVVLRQAALVKRQLRSTDELDVKKDIPIPGTLGLLSEAYDRCSEVCAEYAKTFYLGTMLMTPERRKAIWAIYVWCRRTDELVDGPNASHITPAALDRWEDRLEDVFSGRPFDMLDAALSDTVSKFPVDIQPFRDMIEGMRMDLRKSRYRNFDELYLYCYYVAGTVGLMSVPIMGIAPESKATTESVYNAALALGIANQLTNILRDVGEDARRGRVYLPQDELAQAGLSDEDIFAGRVTDKWRIFMKKQIQRARKFFDEAEKGVTELSAASRWPVLASLLLYRRILDEIEANDYNNFTKRAYVSKPKKLIALPIAYAKSLVPSTRT |
| *Capsicum annuum* Phytoene synthase-1_NCBI_OrioleOrgange_(GU085278) | MSVALLWVVSPCDVSNGTGFLVSVREGNRIFDSSGRRNLACNERIKRGGGKQRWSFGSYLGGAQTGSGRKFSVRSAIVATPAGEMTMSSERMVYDVVLRQAALVKRQLRSTDELDVKKDIPIPGTLGLLSEAYDRCSEVCAEYAKTFYLGTMLMTPERRKAIWAIYVWCRRTDELVDGPNASHITPAALDRWEDRLEDVFSGRPFDMLDAALSDTVSKFPVDIQPFRDMIEGMRMDLRKSRYRNFDELYLYCYYVAGTVGLMSVPIMGIAPESKATTESVYNAALALGIANQLTNILRDVGEDARRGRVYLPQDELAQAGLSDEDIFAGRVTDKWRIFMKKQIQRARKFFDEAEKGVTELSAASRWPVLASLLLYRRILDEIEANDYNNFTKRAYVSKPKKLIALPIAYAKSLVPSTRT |
| *Capsicum annuum* Phytoene synthase-1  ValenciaOrange  (GU085273) | MSVALLWVVSPCDVSNGTGFLVSVREGNRIFDSSGRRNLACNERIKRGGGKQRWSFGSYLGGAQTGSGRKFSVRSAIVATPAGEMTMSSERMVYDVVLRQAALVKRQLRSTDELDVKKDIPIPGTLGLLSEAYDRCSEVCAEYAKTFYLGTMLMTPERRKAIWAIYVWCRRTDELVDGPNASHITPAALDRWEDRLEDVFSGRPFDMLDAALSDTVSKFPVDIQPFRDMIEGMRMDLRKSRYRNFDELYLYCYYVAGTVGLMSVPIMGIAPESKATTESVYNAALALGIANQLTNILRDVGEDARRGRVYLPQDELAQAGLSDEDIFAGRVTDKWRIFMKKQIQRARKFFDEAEKGVTELSAASRWPVLASLLLYRRILDEIEANDYNNFTKRAYVSKPKKLIALPIAYAKSLVPSTRT |
| *Capsicum annuum* Phytoene synthase-1  CM334  (CA04g04080) | MSVALLWVVSPCDVSNGTGFLVSVREGNRIFDSSGRRNLACNERIKRGGGKQRWSFGSCLGGAQTGSGRKFSVRSAIVATPAGEMTMSSERMVYDVVLRQAALVKRQLRSTDELDVKKDIPIPGTLGLLSEAYDRCSEVCAEYAKTFYLGTMLMTPERRKAIWAIYVWCRRTDELVDGPNASHITPAALDRWEDRLEDVFSGRPFDMLDAALSDTVSKFPVDIQPFRDMIEGMRMDLRKSRYRNFDELYLYCYYVAGTVGLMSVPIMGIAPESKATTESVYNAALALGIANQLTNILRDVGEDARRGRVYLPQDELAQAGLSDEDIFAGRVTDKWRIFMKKQIQRARKFFDEAEKGVTELSAASRWPVLASLLLYRRILDEIEANDYNNFTKRAYVSKPKKLIALPIAYAKSLVPSTRT |
| *Capsicum annuum* Phytoene synthase-1  Zunla  (Capana04g002519) | MSVALLWVVSPCDVSNGTGFLVSVREGNRIFDSSGRRNLACNERIKRGGGKQRWSFGSCLGGAQTGSGRKFSVRSAIVATPAGEMTMSSERMVYDVVLRQAALVKRQLRSTDELDVKKDIPIPGTLGLLSEAYDRCSEVCAEYAKTFYLGTMLMTPERRKAIWAIYVWCRRTDELVDGPNASHITPAALDRWEDRLEDVFSGRPFDMLDAALSDTVSKFPVDIQPFRDMIEGMRMDLRKSRYRNFDELYLYCYYVAGTVGLMSVPIMGIAPESKATTESVYNAALALGIANQLTNILRDVGEDARRGRVYLPQDELAQAGLSDEDIFAGRVTDKWRIFMKKQIQRARKFFDEAEKGVTELSAASRWPVLASLLLYRRILDEIEANDYNNFTKRAYVSKPKKLIALPIAYAKSLVPSTRT |
| *Capsicum annuum* Phytoene synthase-1  var. glabriusculum  (Capang04g002191) | MSVALLWVVSPCDVSNGTGFLVSVREGNRIFDSSGRRNLACNERIKRGGGKQRWSFGSCLGGAQTGSGRKFSVRSAIVATPAGEMTMSSERMVYDVVLRQAALVKRQLRSTDELDVKKDIPIPGTLGLLSEAYDRCSEVCAEYAKTFYLGTMLMTPERRKAIWAIYVWCRRTDELVDGPNASHITPAALDRWEDRLEDVFSGRPFDMLDAALSDTVSKFPVDIQPFRDMIEGMRMDLRKSRYRNFDELYLYCYYVAGTVGLMSVPIMGIAPESKATTESVYNAALALGIANQLTNILRDVGEDARRGRVYLPQDELAQAGLSDEDIFAGRVTDKWRIFMKKQIQRARKFFDEAEKGVTELSAASRWPVLASLLLYRRILDEIEANDYNNFTKRAYVSKPKKLIALPIAYAKSLVPSTRT |
| *Capsicum annuum* Phytoene synthase-2  R3  (KX588715) | MSVALLWVVSPNSEVSNGTGFLDSVREGNRVPGRDRNSMWKERIKKGGRQRWNFGSLNAGLRYSDLGGSRTGNGSSFAVQSSLVASPAGEMAVSSEKKVYDVVLKQAALVKRQLRSTDDLEVKPDIVLPGNLGLLSEAYDRCGEVCAESAKTFYLGTLLMTPDRRRAIWAIYVWCRRTDELVDGPNASHITPQALDRWEARLEDIFSGRPFDMLDAALSDTVSRFPVDIQPFRDMVEGMRMDLWKSRYMNFDELYLHCYYVAGTVGLMSVPIMGIAPESKATTESVYNAALALGIANQLTNILRDVGEDARRGRIYLPQDELAQAGLSGEDIFAGRVTDKWRIFMKKQIQRARKFFDQAEKGVAELSSASRWP |
| *Capsicum annuum* Phytoene synthase-2  R7  (KX588716) | MSVALLWVVSPNSEVSNGTGFLDSVREGNRVPGRDRNSMWKERIKKGGRQRWNFGSLNAGLRYSDLGGSRTGNGSSFAVQSSLVASPAGEMAVSSEKKVYDVVLKQAALVKRQLRSTDDLEVKPDIVLPGNLGLLSEAYDRCGEVCAEYAKTFYLGTLLMTPDRRRAIWAIYVWCRRTDELVDGPNASHITPQALDRWEARLEDIFSGRPFDMLDAALSDTVSRFPVDIQPFRDMVEGMRMDLWKSRYMNFDELYLYCYYVAGTVGLMSVPIMGIAPESKATTESVYNAALALGIANQLTNILRDVGEDARRGRIYLPQDELAQAGLSGEDIFAGRVTDKWRIFMKKQIQRARKFFDQAEKGVAELSSASRWPVLASLLLYRKILDEIEANDYNNFTRRAYVSKPKKLLTLPIAYARSLVPPKLTSSSLTKT |
| *Capsicum annuum* Phytoene synthase-2  CM334  (CA02g20350) | MSVALLWVVSPNSEVSNGTGFLDSVREGNRVPGRDRNSMWKERIKKGGRQRWNFGSLNAGLRYSDLGGSRTGNGSSFAVQSSLVASPAGEMAVSSEKKVYDVVLKQAALVKRQLRSTDDLEVKPDIVLPGNLGLLSEAYDRCGEVCAEYAKTFYLGTLLMTPDRRRAIWAIYVWCRRTDELVDGPNASHITPQALDRWEARLEDIFSGRPFDMLDAALSDTVSRFPVDIQPFRDMVEGMRMDLWKSRYMNFDELYLYCYYVAGTVGLMSVPIMGIAPESKATTESVYNAALALGIANQLTNILRDVGEDARRGRIYLPQDELAQAGLSGEDIFAGRVTDKWRIFMKKQIQRARKFFDQAEKGVAELSSASRWPVLASLLLYRKILDEIEANDYNNFTRRAYVSKPKKLLTLPIAYARSLVPPKLTSSSLTKT |
| *Capsicum annuum* Phytoene synthase-2  Zunla  (Capana02g002284) | MSVALLWVVSPNSEVSNGTGFLDSVREGNRVPGRDRNSMWKERIKKGGRQRWNFGSLNAGLRYSDLGGSRTGNGSSFAVQSSLVASPAGEMAVSSEKKVYDVVLKQAALVKRQLRSTDDLEVKPDIVLPGNLGLLSEAYDRCGEVCAEYAKTFYLGTLLMTPDRRRAIWAIYGDVSTFKTYVWCRRTDELVDGPNASHITPQALDRWEARLEDIFSGRPFDMLDAALSDTVSRFPVDIQPFRDMVEGMRMDLWKSRYMNFDELYLYCYYVAGTVGLMSVPIMGIAPESKATTESVYNAALALGIANQLTNILRDVGEDARRGRIYLPQDELAQAGLSGEDIFAGRVTDKWRIFMKKQIQRARKFFDQAEKGVAELSSASRWPVLASLLLYRKILDEIEANDYNNFTRRAYVSKPKKLLTLPIAYARSLVPPKLTSSSLTKT |
| *Capsicum annuum* Phytoene synthase-2  var. glabriusculum  (Capang02g002047) | MSVALLWVVSPNSEVSNGTGFLDSVREGNRVPGRDRNSMWKERIKKGGRQRWNFGSLNAGLRYSDLGGSRTGNGSSFSVQSSLVASPAGEMAVSSEKKVYDVVLKQAALVKRQLRSTDDLEVKPDIVLPGNMGLLSEAYDRCGEVCAEYAKTFYLGTLLMTPDRRRAIWAIYGDVSTFKTYVWCRRTDELVDGPNASHITPQALDRWEARLEDIFSGRPFDMLDAALSDTVSRFPVDIQPFRDMVEGMRMDLWKSRYMNFDELYLYCYYVAGTVGLMSVPIMGIAPESKATTESVYNAALALGIANQLTNILRDVGEDARRGRIYLPQDELAQAGLSGEDIFAGRVTDKWRIFMKKQIQRARKFFDQAEKGVAELSSASRWPVLASLLLYRKILDEIEANDYNNFTRRAYVSKPKKLLTLPIAYARSLVPPKLTSSSLTKT |
| *Capsicum annuum* Phyotene synthase-3  CM334  (CA01g12040) | MRRKLLIRAEVLTVSPKKKEKSIDELLSVQGIAYTHRRIREVVWKQTHVVNDLLCCRNPSFDPVFLDEAYELCRKICAEYAKTFYLGTKLMTEERQKAIWAIYVWCRRTDELVDGPNADYMNNSVLDRWEERLEDIFKNKPYDMLDAALTDTICKFPLDIKPFKDMIDGMRMDTRKSRYANFQELYMYCYCVAGTVGLMSVPIMGIAPECPVSAQTVYNAALHLGIGNQLTNILRDVGEDALRGRVYLPQDELAQYGIRDEDVFARNVTDQWRGFMKEQIRRARFYFNLAEEGASHLNKASRWPVWSSLILYRKILDAIEENDYDNLTKRAYVGRAKKLATLPVSYARALSLPSLAIQ |
| *Aradidopsis*  Phytoene synthase  (NP_197225.1) | MSSSVAVLWVATSSLNPDPMNNCGLVRVLESSRLFSPCQNQRLNKGKKKQIPTWSSSFVRNRSRRIGVVSSSLVASPSGEIALSSEEKVYNVVLKQAALVNKQLRSSSYDLDVKKPQDVVLPGSLSLLGEAYDRCGEVCAEYAKTFYLGTLLMTPERRKAIWAIYVWCRRTDELVDGPNASHITPMALDRWEARLEDLFRGRPFDMLDAALADTVARYPVDIQPFRDMIEGMRMDLKKSRYQNFDDLYLYCYYVAGTVGLMSVPVMGIDPKSKATTESVYNAALALGIANQLTNILRDVGEDARRGRVYLPQDELAQAGLSDEDIFAGKVTDKWRNFMKMQLKRARMFFDEAEKGVTELSAASRWPVWASLLLYRRILDEIEANDYNNFTKRAYVGKVKKIAALPLAYAKSVLKTSSSRLSI |
| *Solanum melongena*  Phytoene synthase  (Sme2.5_00172.1_g00025.1) | MSVALLWVVSPYTEVSNGTGFLDSVREGNSMWNRSSMWKGRFKKGGRQRWNFESLNADSRCSCLGRSRIENGRTFSVQSSLVASPTGEMAVSSEKKVYEVVLKQAALVKRQLVSTDDLEVKPDIVLPGNLGLLSEAYDRCGEVCAEYAKTFYLVWCRRTDELVDGPNASHITPQALDRWEARLEDIFNGRPFDMLDAALSDTVSRFPVDIQPFRDMIEGMRMDLWKSRYKNFDELYLYCYYVAGTVGLMSVPIMGIAPESQATTESVYNAALALGIANQLTNILRDVGEDARRGRVYLPQDELAQAGLSDEDIFAGRVTDKWRIFMKKQIQRARKFFDEAEKGVTELSSASRWPVLASLLLYRKILDEIEANDYNNFTRRAYVSKPKKLLTLPIAYARSLMPPKTTSSPLAKT |
| *Solanum melongena*  Phytoene synthase  (Sme2.5_00721.1_g00007.1) | MSVALLGVVSSCEVSNGTGFLESVREGNRIFDSSRHRNLVSNERIKRGGGKQRWSFGGAQTDNGRKFSVRSAIVATPAGERMMTSEQMVYDVVMRQAALVKRQLRSTDELEVKTDIPIPGNLGLLSEAYDRCGEVCAEYAKTFNLGTMLMTPERRRAIWAIYGEASSHLNIRYVHKQPLYLYEVVVRRTDELVDGPNASYITPTALDRWEDRLENVFNGRPFDMLDAALSDTVSNFPVDIQPFRDMIEGMRMDLRKSRYNNFDELYLYCYYVAGTVGLMSVPIMGIAPESKATTESVYNAALALGIANQLTNILRDVGEDARRGRVYLPQDELAQAGLSDEDIFAGRVTDKWRIFMKKQIQRARKFFDEAEKGVTELTAASRFPVWASLVLYNYAHDSREKKGYLGNIR |
| *Bacillus*  PSY  (WP_013058559.1) | MSVPNKLRDNAMVMLKETSRTFFIPISHLPAELQDAVGSAYLCMRAIDEIEDHPELEAGVKSRLLYAISDLLKKSFNEDEYMALVGPYQADLPDVTLQLGNWIALCPTGVADQVLDATSIMAKGMADWVEKDWHIQNEADLDDYTFYVAGLVGVMLNDIWKWYDGTETDKELAIAFGRGLQSVNILRNTSEDSERGVSFFPNNWSREDMFAYARRNLSLGKKYLEDVKSLPILHFCKIPLALADGTLNALMKGKEKMTRDDVNKTVNEVVDM |
| *Musa cavendishii*  PSY1  (AFP33591.1) | MACLLLRMIAPADIPAGLGSAEAIREGDRFPRKAFRPRNNRNLSRKRRRWSLRSPHADSKYASLRFDPESGMNLPLVSSLLTSTAGEVAVSAEQKVYNVVLKQAALVKQQPRSSTALDVKPDTVIPGSVGLLKEAYDRCGEVCAEYAKTFYLGTLLMTPERRRAIWAIYVWCRRTDELVDGPNASQITPTALDRWESRLDDVFAGRPYDMLDAALSDTVSKYPVDIQPFRDMIEGMRMDLKKSRYKNFDELYLYCYYVAGTVGLMSVPVMGIAPESKATTESVYGAALALGLANQLTNILRDVGEDARRGRIYLPQDELAQAGLSDEDIFGGKVTEKWRSFMKNQIKRARMLFQQAEAGVTELNRASRWPVWASLQLYRQILDEIEANDYNNFTKRAYVSKAKKLLALPVAYGKSLISPSSLSQSNLAKT |
| *Musa cavendishii*  PSY2a  (AFP33592.1) | MSGSVVWVVSPKETSRSSGFSQFAGVKRRWWRSSSAGWSCARSASTARRSVSASLVVTPPRSSEALVYDVVLRQAALVGEARRKRAVDAVEAPPAPLRGDLLDAAYERCGEVCAEYAKTFYLGTMLMTPERRRAIWAIYVWCRRTDELVDGPNASHITPTALDRWSNRLEDLFAGRPYDMYDAALSDTVSNFPVDMQPFKDMIEGMRMDLRKSRYKNFDELYLYCYYVAGTVGLMSVPVMGIAPDSKASAESVYSAALALGIANQLTNILRDVGEDSRRGRVYLPQDELAHAGLSDDDVFEGRVTDKWRTFMKGQITRARMFFDEAEKGIYELNSASRWPVLASLLLYRQILDAIEANDYNNFTKRAYVGKAKKLASLPIAYAKAVIVGPSRFAGTL |
| *Manihot esculenta*  PSY1  (ACY42666.1) | MTIALLWVAIPSTEVSNSFGFFHSVGALDSAKFGSVDRSLMFKKRAKKGMNQKWKSSTVNVDLTNPCIGLGSGSNLPVISSMVASSAGEMAVSSEEKVYNVVLKQAALVKKQLRSSENLDAKTDIAVPGTSSLLSEAYDRCGEVCAEYAKTFYLGTLLMTPERRRAIWAIYVWCRRTDELVDGPNASHITPTALDRWEARLDDVFQGRPFDMLDAALSDTVTKFPVDIQPFKDMIEGMRMDLKKSRYNNFDELYLYCYYVAGTVGLMSVPVMGIAPESQASTESVYNAALALGIANQLTNILRDVGEDARRGRIYLPQDELAQAGLSDEDIFAGEVTNKWRNFMKNQIKRARMFFNEAEKGVTELSAASRWPVWASLLLYKQILDEIEANDYNNFTERAYVNKAKKLAFLPIAYARSFVGSSRVSPPLANP |
| *Manihot esculenta*  PSY2  (ACY42667.1) | MTVALLWVAIPSTEVSNSFGFFHSVRVLDSSKFGCVDRNLMFKGKAKKGRNQKWKSGSVSIDLRSTCIGSGRKLPIISSMVASHAGEIAISSEEKVYNVVLKQAALVKQQLKSSEDLDVKPDIVLPGTLSLLSEAYDRCGEVCAEYAKTFYLGTLLMTPERRRAIWAIYVWCRRTDELVDGPNASHITPTALDRWEARLEDMFRGRPFDMLDAALSDTVTKFPVDIQPFKDMIEGMRMDLKKSRYKNFDELYLYCYYVAGTVGLMSVPVMGIAPESQASTESVYNAALALGIANQLTNILRDVGEDARRGRIYLPQDELAQAGLSDDDIFAGKVTDKWRNFMKNQIKRARMFFNEAEKGVTELSAASRWPVWASLLLYRRILDEIEANDYNNFTKRAYVSKTKKIASLPIAYARSFVGPSRMSSPVTKA |
| *Citrus sinensis*  PSY1-variantX1  (XM_006481880.3) | MSVTLLWVVSPNSQLSNCFGFVDSVREENRLFYSSRFLYQHQTRTAVFNSRPKQFNNSNKQRRNSYPLDTDLRHPCSSGIDLPEISCMVASTAGEVAMSSEEMVYNVVLKQAALVNKQPSGVTRDLDVNPDIALPGTLSLLSEAYDRCGEVCAEYAKTFYLGTLLMTSERRRAIWAIYVWCRRTDELVDGPNASHITPTALDRWESRLEDLFRGRPFDMLDAALSDTVTKFPVDIQPFRDMIEGMRMDLRKSRYKNFDELYLYCYYVAGTVGLMSVPVMGIAPDSQATTESVYNAALALGIANQLTNILRDVGEDARRGRVYLPQDELAQAGLSDDDIFAGEVTIKWRNFMKNQIKRARMFFDMAENGVTELSEASRWPVWASLLLYRQILDEIEANDYNNFTKRAYVSKAKKIAALPIAYAKSLLRPSRIYTSKA |
| *Citrus sinensis*  PSY2  (XM_006492653) | MCSTFSLAAKPSIGESRGKIWSLQVINSAARSELNINIPREKNSIILPRLSTQGMLQSDVHICEIAERQSLANNLSDKGDAFRSKPQLDAMFLDEAYERCRKICAEYAKTYYLGTLLLTKERQKAIWAVYAWCKRTDELVDGPNAFCVSPAALDRWEERLQDIFNGHPNDILDAALTDTVSKFNLHIKPFRDMIKGMRMDTEKCRYANFQELYLYCYYAGGTVGLMSVPVMGMDPDSSASAQSIYNAALYLGIGNQLTNILRDVGEDASRGRIYLPQDELAKFGLRDEDIFSRKVSDEWREFMKEQITRARYFYKLAEEGASELDKASRWPVWTVLILYQKILDAIQDNDYDNLTKRAYVGNIKKLFLVRQAYYRAQSISTATD |
| *Triticum durum*  Phytoene synthase-A1  (EU096090) | MATTVTLLLGAASSPGPAAGDGAARDGFQCSRLLPKKKQQRPRWVLCSLKYGCLGVGEPGEAGGRSAASPVYSSLTVSPGGDAAVAVVSSEQKVYDVVVKQAALLKRQLRPSQQQQQAPPAVARELDAPRGGLGEAYARCGEICEEYAKTFYLGTLLMTEERRRAIWAIYVWCRRTDELVDGPNASHITPQALDRWERRLEDLFAGRPYDMLDAALSDTITKFPIDIQPFKDMIDGMRTDLKKARYKNFDELYMYCYYVAGTVGLMSVPVMGIAPDSKATAETVYGAALALGLANQLTNILRDVGEDARRGRIYLPQDELAEAGLSDEDIFKGVVTDKWRKFMKRQIKRARMFFEEAERGVTELRKESRWPVWASLLLYRQILDEIEANDYNNFTKRAYVGKAKKVLALPVAYGRSLLLPYSLRNNQT |
| *Triticum durum*  Phytoene synthase-B1  (EU096092) | MATTVTLLLGAASSPGDGAARDGVQCSRMLPRRRQQRPRWVLCSLKYGCLGVGEPGEAGTRSAASPVYSSLTVSPGGEAAVAVVSSEQKVYDVVVKQAALLKRQLRPQQAAPPAVARELDAPRGGLGEAYARCGEICEEYAKTFYLGTLLMTEERRRAIWAIYVWCRRTDELVDGPNASHITPQALDRWERRLEDLFAGRPYDMLDAALSDTITKFPIDIQPFKDMIDGMRTDLKKARYTNFDELYMYCYYVAGTVGLMSVPVMGIAPESKATAESVYGAALALGLANQLTNILRDVGEDARRGRIYLPQDELAEAGLSDEDIFKGVVTDKWRKFMKRQIKRARMFFEEAERGVTELRKESRWPVWASLLLYRQILDEIEANDYNNFTRRAYVGKAKKVLALPVAYGRSLLLPYSLRNNQT |
| *Eriobotrya japonica*  Phytoene synthase-1  (AIT18246.1) | MSVALIWVVSANTEVFKFYGILDSSRFVLGNRSSIRAKMGAKQDWKSCSLSTHVKYSSVGGSGLGSETKFPVLLSMVANPLGESAVSSEQKVYDVVLKQASLGKKQLSSNGYLDVKRDIILPGNLSLLSKAYDRCGEVCAEYAKTFYLGTLLMTPERRRAIWAIYVWCRRTDELVDGPNASHITPTALDRWESRLDDLFQGRPFDMLDAALSDTVTKFPVDIQPFKDMIEGMRMDLRKSRYQSFDELYLYCYYVAGTVGLMSVPVMGISPESQATTESVYNAALALGIANQLTNILRDVGEDARRGRIYLPQDELAEAGLSDADIYAGKVTDKWRSFMKDQIKRARMFFDEAEKGVTELSEASRWPVLASLLLYRQILDEIEANDYNNFTRRAYVSKAKKLLALPIAYTKSIIRPPRTSPELRKYNL |
| *Eriobotrya japonica*  Phytoene synthase-2A  (AIT18247.1) | MSVVLLWVVSPKQNASSLLGLMPRICTPRRSKFCPKLGFSSRVLAYSGAVVNPARSSEEKVYEVVLKQAALVEEQSTVKRKSLDLDERIVTEGLDNWDLLDKAYDRCGEVCAEYAKTFYLGTLLMTPERRRAVWAIYVWCRRTDELVDGPNASYITPKALDRWEKRLTDLFEGRPYDMYDAALSDTVTKYPVDIQPFRDMVEGMRLDLRKSRYQNFDELYLYCYYVAGTVGLMSVPVMGICPESRASTESVYNAALALGIANQLTNILRDVGEDARRGRVYLPQDELAQAGLSDDDIFRGKVTDKWQSFMKGQIQRARMFFDEAEKGVSELNSASRWPVWASLLLYRQILDVIEANGYDNFTKRAYVGKAKKFVSLPVAYGRAIIGPSKLTKQLVPR |
| *Eriobotrya japonica*  Phytoene synthase-2B  (AIT18249.1) | MSGVLLWVVSPKENANSLLGLLPRICTPRRSKLCSKLGFSSGVLAYSGAVANPARSSEEKVYEVVLKQAALVREPNTVKKKSLDLDERITDGLNNWDLLNKAYDRCGEVCAEYAKTFYLGTLLMTPERRRAVWAIYVWCRRTDELVDGPNASYITPKALDRWEKRLTDLFEGRPYDMYDAALSDTVAKYPVDIQPFRDMVEGMRLDLRKSRYQNFDELYLYCYYVAGTVGLMSVPVMGISPESKASTESVYNAALALGIANQLTNIPRDVGEDARRGRVYLPQDELAQAGLSDNDIFRGKVTDKWQSFMKGQIKRARMFFDEAEKGVSELNSASRWPVWASLLLYRQILDAIEANGYDNFTKRAYVGKAKKLASLPVAYGRAILGPSKLTKQLVPR |
| *Cucumis melo*  Phytoene synthase-1  (AEH03200.1) | MSLASSLVVSSNVELSPSSFGFLDSVRDGPQIPDSFRFSSRNRVPNLINKKQKWGNHSHSTELKYPILHESGYGSVIVASMVANPAGEIAVSAEQKVYNVVMKQAALVKRQLRTAGELDVKPDIVLPGTLSLLNEAYDRCGEVCAEYAKTFYLGTMLMTPERQKAIWAIYVWCRRTDELVDGPNASHITPTALDRWEARLEELFQGRPFDMLDAALADTVTKFPVDIQPFKDMIEGMRMDLRKSRYKNFDELYLYCYYVAGTVGLMSVPVMGIAPESQASTESVYNAALALGIANQAPPNILRDVGEDARRGRIYLPQDELAQAGLSDEDIFAGRVTDKWRNFMKNQIKRARMFFDEAEKGVLELNKASRWPVWASLLLYRQILDEIEANDYDNFTKRAYVSKAKKILALPMAYGRALLGPS |
| *Cucumis melo*  Phytoene synthase-2  (AEH03199.1) | MSSCVFVTPKSLTFIGECKGNVFPKRFKTIRNGGGIIAAPRSSETLKLQTLSKQGIPLDHLKVEEIVERQSEGNHFLREEWCKKKMKFQPSFLEEAYESCRKICAEYAKTFYLGTLLMTKERQRAIWAIYVWCRRTDELVDGPNAVYMNPKVLDRWEERLEDIFEGRPYDLLDAALSHTVSKFPIDMKPFRDMIEGMRMDTKKCRYENFEELYLYCYYVAGTVGLMSVPVMGIAPDSSLSTHTIYSAALHLGIGNQLTNILRDVGEDATRGRVYLPQDELAQFGLCDADILRMRVSDKWREFMKEQIKRARFYFKLAEEGASQLDKASRWPVWSSLMLYRKILEAIEENDYNNFTKRAYVGRSKKLLALPLAYTKSISPPTLVFN |
| *Solanum tuberosum*  Phytoene synthase-1  (Sotub03g008370.1.1) | MSVALLWVVSPCEVSNGTGFLESVREGKSFFDSSRHRNLVSNERINRGGGKQTNNGRKFSVRSAIVATPSGERTMTSEQMVYDVVLRQAALVKRQLRSTDELEVKPDIPVPGNLGLLSEAYDRCGEVCAEYAKTFNLGTMLMTPERRRAIWAIYVWCRRTDELVDGPNASYITPAALDRWEDRLEDVFNGRPFDMLDGALSDTVSNFPVDIQPFRDMIEGMRMDLRKSRYKNFDELYLYCYYVAGTVGLMSVPIMGIAPESKATTESVYNAALALGIANQLTNILRDVGEDARRGRVYLPQDELAQAGLSDEDIFAGRVTDKWRIFMKKQIHRARKFFDDAEKGVTELSAASRFPVWASLVLYRKILDEIEANDYNNFTRRAYVSKSKKLIALPIAYAKSLVPPTRTISLLS |
| *Solanum tuberosum* Phytoene synthase-2  (Sotub02g024440.1.1) | MSVALLWVVSPNSEVLNGTGFLDSVREGNRGLESSRFPSPNRNSMWKGRFKKGGRQEWNFGFLNADLRYSCLGRSRTENGRSFSVQSSLVASPAGEMAVSSEKKVYEVVLKQAALVKRHLISTEDIEVKPDIVVPGNLGLLSEAYDRCGEVCAEYAKTFYLGTMLMTPDRRRAIWAIYVWCRRTDELVDGPNASHITPQALDRWEARLEDIFNGRPFDMLDAALSDTVSKFPVDIQPFRDMVEGMRMDLWKSRYNNFDELYLYCYYVAGTVGLMSVPIMGIAPESKATTESVYNAALALGIANQLTNILRDVGEDARRGRVYLPQDELAQAGLSDEDIFAGRVTDKWRIFMKKQIQRARKFFDEAEKGVTELSSASRWPVLASLLLYRKILDEIEANDYNNFTRRAYVSKPKKLLTLPIAYARSLVPPKSTSSPLAKT |
| *Oryza sativa*  Phytoene synthase-1  (AAS18307.1) | MAAITLLRSASLPGLSDALARDAAAVQHVCSSCLPSNNKEKKRRWILCSLKYACLGVDPAPGEIARTSPVYSSLTVTPAGEAVISSEQKVYDVVLKQAALLKRHLRPQPHTIPIVPKDLDLPRNGLKQAYHRCGEICEEYAKTFYLGTMLMTEDRRRAIWAIYVWCRRTDELVDGPNASHITPSALDRWEKRLDDLFTGRPYDMLDAALSDTISKSPIDIQPFRDMIEGMRSDLRKTRYKNFDELYMYCYYVAGTVGLMSVPVMGIAPESKATTESVYSAALALGIANQLTNILRDVGEDARRGRIYLPQDELAEAGLSDEDIFNGVVTNKWRSFMKRQIKRARMFFEEAERGVTELSQASRWPVWASLLLYRQILDEIEANDYNNFTKRTYVGKAKKLLALPVAYGRSLLMPYSLRNSQK |
| *Oryza sativa*  Phytoene synthase-2  (AAK07735.1) | MTGEGSPNQNCRGAPRGLLAGGFEGGPPPPRQVEQDSSHFSQGYNEGEEAGCCRLVGEAAAGREPGGGHGAGGRGVEEDGGGEASRGRLGTRCGRPHRAARGRRGGRGGRVDWGLLLGDAYHRCGEVCAEYAKTFYLGTQLMTPERRKAVWAIYVWCRRTDELVDGPNSSYITPKALDRWEKRLEDLFEGRPYDMYDAALSDTVSKFPVDIQPFKDMIEGMRLDLWKSRYRSFDELYLYCYYVAGTVGLMTVPVMGIAPDSKASTESVYNAALALGIANQLTNILRDVGEDSRRGRIYLPLDELAEAGLTEEDIFRGKVTDKWRKFMKGQILRARLFFDEAEKGVAHLDSASRWPVLASLWLYRQILDAIEANDYNNFTKRAYVNKAKKLLSLPVAYARAAVAS |
| *Oryza sativa*  Phytoene synthase-3  (ABC75828.1) | MAPPPPPPCSVRAAGSNPIGCLEVAEPWSGAAPPPLPPLPGHLHVAAPAAEDDDDALAAAAAAVPSEQRVHDVVLKQAALAAAAPEMRRPAQLAERERVAGGLNAAFDRCGEVCKEYAKTFYLATQLMTPERRRAIWAIYVWCRRTDELVDGPNASHMSALALDRWESRLDDIFAGRPYDMLDAALSHTVATFPVDIQPFRDMIEGMRLDLTKSRYRSFDELYLYCYYVAGTVGLMTVPVMGISPDSRANTETVYKGALALGLANQLTNILRDVGEDARRGRIYLPMDELEMAGLSEDDIFDGRVTDRWRCFMRDQITRARAFFRQAEEGASELNQESRWPVWASLLLYRQILDEIEANDYNNFTKRAYVPKAKKIVALPKAYYRSLMLPSSVRHCSSLTSS |
| *Crocus sativus*  Phytoene synthase-1a  (QCI61139.1) | MAIALLRVFSPIEFPIGLGVEGDLRGKVVTKYDGNLERKKQRWKMCSLYTDSKYACVSFDPEGNGNFMIRSSMVASPREEFISSEQRVYDVVLKQAALVREQGRLTTPLDVKPDMAVPGALYLLKEAYDRCGEVCAEYAKTFYLGTLLMTPERRRAIWAIYVWCRRTDELVDGPNASHITPSALDRWESRLEDLFAGRPYDMFDAALSDTVSQFPVDIQPFKDMVEGMRLDLKKSRYKNFDELYLYCYYVAGTVGLMSVPVMGIAPESQATTESVYNAALSLGIANQLTNILRDVGEDARRGRIYLPQDELAQAGLSDEDVFSGKVTDKWRNFMKKQIKRARMFFQEAEKGVTELSQASRWPVLASLLLYRQILDEIEKNDYNNFTKRAYVSKAKKLASLPIAYGRSLISPASIKRQSSLLKG |
| *Crocus sativus*  Phytoene synthase-1b  (QCI61140.1) | MATQMMRVVSPAEVSVVVGHSRLSPMGSSRRKSVGKCSVAPSLVANPVKDVAMSSEQKVYDVILKQAALVEQQLRSKSNVSLDVNLELVIQGAPYLSKDAYDRCGQVCAEYAKTFYLGTLLMTPERRKAIWAIYVWCRRTDELVDGPNASYITPSALDRWEAKLEDLFAGRPYDMFDAALSDTVSKFPVDIQPFKDMIKGMRMDLKKSRYKNFDELYLYCYYVAGTVALMSVPVMGIAPDSQATTESVYDAALALGIANQLTNILRDVGEDARRGRIYLPQDELAQSGLSDEDVFNGKVTDKWRNFMKSQIRRAREFFLEAEKGVTELSLASRWPVLASLLLYREILDEIEANDYNNFTKRAYVGKAKKIMALPVAFGRSLVSPLRRKQGSQGNA |
| *Crocus sativus*  Phytoene synthase-2  (QCI61141.1) | MSGSVVLLISSGETTMTSTFPHHFTGRNNVNGKVRKRVVRIMPLIPSKGFTVSSISVTTDKPSSSVDIVYEVIVKQAKLVREQKVRASSVGPKDRNLLNEAYDRCQEVCAEYAKTFYLGTLLMTPERQRAIWAIYVWCRRTDELVDGPNASYITPTALDRWEKRLNDLFEGHPYDLYDDALSDTVNKFPIDIQPFRDMIEGMRSDLKKSRYQNFDELYLYCYYVAGTVGLMSVPVMGISPYSKASAEEVYNSALALGIANQLTNILRDVGEDARRGRIYLPQDELAQAGLSDDDIFRGRVTDKWRNFMKGQIRRARMFFDEAEKGVTELNSASRWPVLASLLLYRQILDAIEVNDYNNFTKRAYVGKTKKLVSLPLAYARALAGPSKV |
| *Crocus sativus*  Phytoene synthase-3  (QCI61142.1) | MSTSILPNLSMNYSNPIRCTIQLPSIPRHDKISNIPTKLPKSHGISLTELQVNEVVVRQSRCLPTSDGNRKRVSLNATFLEEGYRRCRGVCEEYAKTFYLGTQLMTEERQRAIWAIYVWCRRTDELVDGPNSVHTNKVVLDRWQERLEDIFQGRPYDMFDAALTDTIFKFPIDIKPFKDMIEGMRMDTNKWRYLSFQELYLYCYYVAGTVSLMSVPVMGIAPESHTPIHRVYEAAVDLGIGMQLTNILRDVGEDASRNRIYLPQDELAQFGLSDEDIFSRKVTSRWREFMTEQISRARLYFDQAEEGASQLNEASRWPVWASLLLYREILDSIEDNDYDNLTKRAYVGRAKKFLMLPWGFAKAVNL |
| *Sorghum bicolor*  Phytoene synthase-1  (AAW28996.1) | MAIILVRAASPAGLSDAAGIISHHGSLQCSSLPLLNKRPAPARRWMLCSLRYGCLGLDPGRPSPPAVYSSLAVNPAGEAVVSSEQKVYDVVLKQSALLKRQLRKPVLDVVRPQEDLAEMPRNGLNQAYDRCGEICEEYAKTFYLGTMLMTEERRRAIWAIYVWCRRTDELVDGPNANYITPTALDRWEKRLEDLFAGRPYDMLDAALSDTISRFPIDIQPFRDMIEGMRSDLRKTRYNNFDELYMYCYYVAGTVGLMSVPVMGIAPESKATTESVYSAALALGIANQLTNILRDVGEDATRGRIYLPQDELAQAGLSDEDIFKGVVTNRWRNFMKRQIKRARMFFEEAERGVTELSQASRWPVWASLLLYRQILDEIEANDYNNFTKRAYVGKGKKLLALPVAYGKSLLLPCSLRNTQT |
| *Sorghum bicolor*  Phytoene synthase-3  (AAW28997.1) | MMSTSHAVKQSPACAARWRRRQHGQRSADAPARTATFSARHGRQRRGGRGVAPCSVRAAGSNTIGCLEAEAWGGAAHAPGSAPSLLLPSFHVEAPAPGGDALAVPSEQRVQEVVLKQAALAAAAPRTARIEPVPMDGGLKAAFHRCGEVCQEYAKTFYLATQLMTPERRRAIWAIYVWCRRTDELVDGPNASHISAVALDRWESRLEDIFAGRPYDMLDAALSDTVANFPVDIQPFRDMIEGMRMDLKKSRYRSFDELYLYCYYVAGTVGLMSVPVMGISPDSRAATETVYKGALALGLANQLTNILRDVGEDARRGRIYLPQDELEMAGLSDADILDGRVTDEWRSFMRGQITRARAFFRQAEEGATELNQESRWPVWASLLLYRQILDEIEANDYNNFTRRAYVPKTKKLMALPKAYLRSLVPPSSSQAERQRHHSSLT |
| *Solanum lycopersicum*  Phytoene synthase-1  (Solyc03g031860) | MSVALLWVVSPCDVSNGTSFMESVREGNRFFDSSRHRNLVSNERINRGGGKQTNNGRKFSVRSAILATPSGERTMTSEQMVYDVVLRQAALVKRQLRSTNELEVKPDIPIPGNLGLLSEAYDRCGEVCAEYAKTFNLGTMLMTPERRRAIWAIYVWCRRTDELVDGPNASYITPAALDRWENRLEDVFNGRPFDMLDGALSDTVSNFPVDIQPFRDMIEGMRMDLRKSRYKNFDELYLYCYYVAGTVGLMSVPIMGIAPESKATTESVYNAALALGIANQLTNILRDVGEDARRGRVYLPQDELAQAGLSDEDIFAGRVTDKWRIFMKKQIHRARKFFDEAEKGVTELSSASRFPVWASLVLYRKILDEIEANDYNNFTKRAYVSKSKKLIALPIAYAKSLVPPTKTASLQR |
| *Solanum lycopersicum*  Phytoene synthase-2  (Solyc02g081330) | MSVALLWVVSPNSEVSYGTGFLDSVREGNRGLESSRFPSRDRNSMWKGGFKKGGRQGWNFGFLNADLRYSCLGRSRTENGRSFSVQSSLVASPAGEMAVSSEKKVYEVVLKQAALVKRHLISTDDIQVKPDIVLPGNLGLLSEAYDRCGEVCAEYAKTFYLGTMLMTPDRRRAIWAIYVWCRRTDELVDGPNASHITPQALDRWEARLEDIFNGRPFDMLDAALSDTVSRFPVDIQPFRDMVEGMRMDLWKSRYNNFDELYLYCYYVAGTVGLMSVPIMGIAPESKATTESVYNAALALGIANQLTNILRDVGEDARRGRVYLPQDELAQAGLSDEDIFAGKVTDKWRIFMKKQIQRARKFFDEAEKGVTELSSASRWPVLASLLLYRKILDEIEANDYNNFTRRAYVSKPKKLLTLPIAYARSLVPPKSTSSPLAKT |
| *Citrullus lanatus*  PSY  (AGT57744) | MSFAPSLVVSPNVELSPSSFGFLDSVRDGSQIPDSSRFFSRNRAANLMSKKQKWGNHSHSTELKYPILCEGGYGSVIAASMVANPAGEMAVSAEQKVYNVVMKQAALVKRQLRTAGELDVKPDIVLPGTLSLLNEAYDRCGEVCAEYAKTFYLGTMLMTPERQKAIWAIYVWCRRTDELVDGPNASHITPTALDRWEARLEELFQGRPFDMLDAALADTVTKFPVDIQPFKDMIEGMRMDLRKSRYKNFDELYLYCYYVAGTVGLMSVPIMGIAPDSQASTESVYNSALALGIANQLTNILRDVGEDARRGRIYLPQDELAQAGLSDEDIFAGRVTDKWRNFMKSQIKRARMFFDEAEKGVLELNKASRWPVWASLLLYRQILDEIEANDYNNFTKRAYVSKAKKILALPMAYARSLLGPS |
| *Zea Mays*  Phytoene synthase-1  (AAR08445.1) | MAIILVRAASPGLSAADSISHQGTLQCSTLLKTKRPAARRWMPCSLLGLHPWEAGRPSPAVYSSLAVNPAGEAVVSSEQKVYDVVLKQAALLKRQLRTPVLDARPQDMDMPRNGLKEAYDRCGEICEEYAKTFYLGTMLMTEERRRAIWAIYVWCRRTDELVDGPNANYITPTALDRWEKRLEDLFTGRPYDMLDAALSDTISRFPIDIQPFRDMIEGMRSDLRKTRYNNFDELYMYCYYVAGTVGLMSVPVMGIATESKATTESVYSAALALGIANQLTNILRDVGEDARRGRIYLPQDELAQAGLSDEDIFKGVVTNRWRNFMKRQIKRARMFFEEAERGVTELSQASRWPVWASLLLYRQILDEIEANDYNNFTKRAYVGKGKKLLALPVAYGKSLLLPCSLRNGQT |
| *Zea Mays*  Phytoene synthase-2  (AAX13807.1) | MAAGSSAVWAAQHPACSGGKFHHLSPSHSHCRPRRALQTPPALPARRSGASPPRASLAAAAPAVAVRTASEEAVYEVVLRQAALVEAATPQRRRTRQPRWAEEEEEERVLGWGLLGDAYDRCGEVCAEYAKTFYLGTQLMTPERRKAVWAIYVWCRRTDELVDGPNASYITPTALDRWEKRLEDLFEGRPYDMYDAALSDTVSKFPVDIQPFKDMVQGMRLDLWKSRYMTFDELYLYCYYVAGTVGLMTVPVMGIAPDSKASTESVYNAALALGIANQLTNILRDVGEDARRGRIYLPLDELAQAGLTEEDIFRGKVTGKWRRFMKGQIQRARLFFDEAEKGVTHLDSASRWPVLASLWLYRQILDAIEANDYNNFTKRAYVGKAKKLLSLPLAYARAAVAP |
| *Zea Mays*  Phytoene synthase-3  (ABC75827.1) | MMSTSRAVKSPACAARRRQWSADAPNRTATFLACRHGRRLGGGGGAPCSVRAEGSNTIVCLEAEAWGGAPALPGLRVAAPSPGDAFVVPSEQRVHEVVLRQAALAAAAPRTARIEPVPLDGGLKAAFHRCGEVCREYAKTFYLATQLMTPERRIAIWAIYVWCRRTDELVDGPNASHISALALDRWESRLEDIFAGRPYDMLDAALSDTVARFPVDIQPFRDMIEGMRMDLKKSRYRSFDELYLYCYYVAGTVGLMSVPVMGISPASRAATETVYKGALALGLANQLTNILRDVGEDARRGRIYLPQDELEMAGLSDADVLDGRVTDEWRGFMRGQIARARAFFRQAEEGATELNQESRWPVWSSLLLYRQILDEIEANDYDNFTRRAYVPKTKKLMALPKAYLRSLVVPSSSSQAESRRRYSTLT |

**Table S5. Putative transcriptional regulatory elements identified in PSY-1; corresponds Fig. S1.**

| Ref | Regulatory element | Regulatory element; binding factor | Gene | Plant | Gene function; Response |
| --- | --- | --- | --- | --- | --- |
| (a) | ATtTTGACtCAT | WRKY28 BS; WRKY28 | ICS1 | *Arabidopsis thaliana* | SA biosynthesis; Biotic stress response (van Verk et al., 2011) |
| (b) | AtAACGTGTTaA | T/G box; JAMYC2 and JAMYC10 | LAP | *Lycopersicon esculentum* | Amino acid hydrolysis; Wounding response (Boter et al., 2004). |
| (c) | AAAAtGTTAAAA | Box I; GT-1 | Cab-E | *Nicotiana plumbaginifolia* | Chlorophyll a binding;  Light regulated (Schindler and Cashmore, 1990) |
| (d) | GaACCCATaAAG | Box E; unknown | CCoAOMT | *Petroselinum crispum* | Phenylpropanoid biosynthesis; Stress response (Grimmig and Matern, 1997). |
| (e) | CGgGTTTGGTCG | Box II; unknown | psCHS1 | *Pisum sativum* | Flavonoid biosynthesis;  Stress response (UV and elicitor) (Ito et al., 1997). |
| (f) | TTTTAACaTTTT | Box I; GT-1 | Cab-E | *Nicotiana plumbaginifolia* | Chlorophyll a binding;  Light regulated (Schindler and Cashmore, 1990). |
| (g) | CTACGTGTA | ACE1;  HY5 | FHY1 | *Arabidopsis thaliana* | Phytochrome a regulation; light regulated (Li et al., 2010). |
|  | CTACGTGT | GBF1 BS7; GBF1 |  | *Arabidopsis thaliana* | Stress regulated |
| (h) | CCAAAATTaG | CarG (016); RIN | TBG4 | *Lycopersicon esculentum* | Cell wall modification;  Ripening regulated (Fujisawa et al., 2011). |
|  | CCAAAATTAG | CarG;  FLC | SOC1 | *Arabidopsis thaliana* | TF;  Flowering (Hepworth et al., 2002). |
|  | TGACTAAATATAGAAA | CarG3 (DTA4) | AGL15 | *Arabidopsis thaliana* | TF;  Plant development (Tang and Perry, 2003). |
| (i) | AGTGTAAAAt | Box III*;  unknown | Cab1R | *Oryza sativa* | Chlorophyll a binding;  Light regulated (Luan and Bogorad, 1992). |
| (j) | TGGTAGGTAag | MRE-core;  R2R3-type MRE | F3H | *Arabidopsis thaliana* | Flavonoid biosynthesis;  Light regulated (Hartmann et al., 2005). |
|  | TGGTAGGTaAGA | Box I; unknown | chsA | Various  plants | Flavonoid biosynthesis;  Light regulated |
|  | GGTAGGTaAGA | Box L;  unknown | PAL-1 | *Petroselinum crispum* | Phenylpropanoid  Biosynthesis;  Unknown (Logemann et al., 1995). |
|  | TGGTAGGTaAGAT | 13bp - box | LTR-Tto1 | *Nicotiana tabacum* | LTR-retrotransposon;  Stress response |
|  | GGTAGGTaAGA | MYB92 BS3;  MYB92 |  | *Glycine max* | Unknown;  Stress response |
|  | TGTCTTGTTtT | MYCS (PT3); MYC | PT3 | *Solanum melongena*  *Nicotiana tabacum* | Mycorrhizal induced (Chen et al., 2011). |

**Table S6. Putative transcriptional regulatory elements identified in PSYS-2 promoter. Corresponds to Fig. S2.**

| Ref | Motif | Regulatory element; binding factor | Gene | Plant | Gene function; Response | In PSY1 |
| --- | --- | --- | --- | --- | --- | --- |
| (a) | CTTAATGGGTCT | Box E | CCoAOMT | *Petroselinum crispum* | Phenylpropanoid biosynthesis; Stress response (Grimmig and Matern, 1997). | Y |
| (b) | AATtTCCAACCA | Box-L4; DcMYB1 | DcPAL1 | *Daucus carota* | Phenylpropanoid biosynthesis; Stress response (Maeda et al., 2005). | N |
| (c) | tGGCGGaCAGTG | GRA; unknown | Rab17 | *Zea mays* | ABA regulated | N |
|  | aTTTACGTGTAt | C/A box; HY5 | PIN1 | *Arabidopsis thaliana* | Auxin carrier proteins; cytokinin regulated | N |
|  | TTACGTGT | GBF1 BS8; GBF1 |  | *Arabidopsis thaliana* | Stress regulated | Y |
| (d) | CCGAAATAAG | CarG; FLC | SOC1 | *Arabidopsis thaliana* | Transcription factor;  Flowering (Hepworth et al., 2002). | Y |
| (e) | TtaTTATTATTATT ATTATa | C2; unknown | GmAux28 | *Glycine max* | Auxin regulated | N |
|  | CTCAAACCgAaCAA | P box 4; unknown | C4H | *Arabidopsis thaliana* | Phenylpropanoid biosynthesis; Stress response (Bell-Lelong et al., 1997). | N |
| (f) | TTCTGACCAG | W-box; bhWRKY1 | bhGOLS1 | *Boea hygrometrica* | Sugar biosynthesis; Stress response (Wang et al., 2009). | N |
|  | AAAAGGTTAAAA | Box I; GT-1 | Cab-E | *Nicotiana plumbaginifolia* | Chlorophyll a binding;  Light regulated (Schindler and Cashmore, 1990). | Y |
| (g) | AATTATAATCA | C-DP BS2; cytokinin dependent protein | POR | *Cucumis sativus* | Plastid development; cytokinin regulated (Fusada et al., 2005). | N |
| (h) | TCTACGTGTA | UV LRE; unknown | CHS | *Arabidopsis thaliana* | Flavonoid biosynthesis;  Light regulated (Wade et al., 2001) | Y |
|  | TTcTACGTGTAA | G/ A box;  FY5 | IAA3/ | *Arabidopsis thaliana* | Auxin regulation; cytokinin regulated | N |
|  | CTACGTGT | ACE1;  FY5 | FHY1 | *Arabidopsis thaliana* | Phytochrome a regulation; light regulated (Li et al., 2010). | Y |
|  | CTACGTGT | GBF1 BS7; GBF1 |  | *Arabidopsis thaliana* | Stress regulated | Y |
|  | TACGTGTAAATC | ABRE3a.3b; unknown | Sbdhn2 | *Sorghum bicolor; Sorghum vulgare* | ABA regulated  (Buchanan et al., 2004). | N |
|  | TaACTATATATAGTA | CarG3 (DTA4) | AGL15 | *Arabidopsis thaliana* | Transcription factor;  Plant development | Y |
| (i) | TTGcTtCTTGTtG  ATGA | Motif d; epicotyl-specific | PSPAL2 | *Pisum sativum* | Phenylproponoid biosynthesis;  Unknown (Kato et al., 1995). | N |
| (j) | CCAAAGTTGG | CarG (016); RIN |  | *Lycopersicon esculentum* | Cell wall modification;  Ripening regulated (Tang and Perry, 2003). | Y |

**Table S7. Putative regulatory elements identified in the DXS promoter region. Table corresponds to Fig. 5.**

| Ref | Motif | Regulatory element; Binding factor | Gene | Plant | Gene function; Response |
| --- | --- | --- | --- | --- | --- |
| (a) | tTCtGACGTGGC | G-box; CG-1 | CAB | *Nicotiana plumbaginifolia* | Chlorophyll binding; light regulated (Schindler and Cashmore, 1990). |
|  | CTgACGTGGC | CUF-1 BS;CUF-1 | CAB2 | *Arabidopsis thaliana* | Chlorophyll binding; light regulated |
|  | TGACGTGGC | TAGA2 | GH3 | *Glycine max* | Auxin regulated |
|  | tGACGTGG | G-box; GBF | gPAL2 | *Antirrhinum majus* | Phenylpropanoid biosynthesis; flowering related (Liang et al., 1989). |
|  | ACGTGGC | ABRE1/2; ABI3; ABI5; AREB1 | RD29B | *Arabidopsis thaliana* | Drought response; ABA regulated (Uno et al., 2000). |
|  | TGACGTGG | GBF1 BS4; GBF |  | *Arabidopsis thaliana* | Stress regulated |
|  | TGACGTGG | Hex; HBP-11(17); HBP-1b(c38) | H3 | *Triticum aestivum* | Histones (Mikami et al., 1994). |
| (b) | TCTCCATTTc | Gap box 1; GAPF | GapA | *Arabidopsis thaliana* | Glycolysis; light regulated |
|  | TGgGtTTATTGTT | CII; S2F | RPL21 | *Spinacia oleracea* | Plastid development; developmental regulated (Lagrange et al., 1997). |
| (c) | ACAgAAAAaAAAAAAA GaaA |  | SBEI | *Zea mays* | Starch branching; Sugar regulated (Kim and Guiltinan, 1999). |
|  | AAAAAAAAAAGAAAAA | UN U2; unknown | rbcS3C | *Lycopersicon esculentum* | Carbon fixation; light regulated (Manzara et al., 1991). |
|  | AAAAAAGAAAGAAA | Box d; DOF1 | cyPPDK1 | *Zea mays* | Photosynthesis;  Photosynthesis regulated (Yanagisawa, 2000). |
| (d) | CCAgCAcACCCC | Box 2; unknown | chsA | *Petunia hybrida* | Phenylpropanoid biosynthesis; flowering regulated (van der Meer et al., 1990). |
| (e) | TGGGTGGTTGGG | Box L3; DcMYB1 | DcPAL1 | *Daucus carota* | Phenylpropanoid biosynthesis; Stress response (Maeda et al., 2005). |
|  | GGGTGGT | GS1B AC box 1; PtMYB1 and 4 | PtGS1b | *Pinus sylvestris* | Nitrogen metabolism (Cánovas et al., 2007). |
